# Chronic Cadmium Exposures and Hyperglycemia Additively Drive Mitochondrial Dysfunction in Hepatic Cells: Key Implications for MASLD Etiopathogenesis

**DOI:** 10.1101/2025.09.13.676023

**Authors:** Rahul Kumar, Ashwin Chinala, Li Chen, Sharina Desai, Marcus A. Garcia, Sarah J. Blossom, Matthew J. Campen, Rama R. Gullapalli

## Abstract

Effects of chronic heavy metal stress on hepatocellular pathophysiology remains ill-understood. Human livers are a long-term accumulative site for many toxic heavy metals (e.g., cadmium and arsenic) whose effects are unknown. In the current study, we studied effects of chronic, low-dose exposures of cadmium (CLEC) modulated by normoglycemic (5.6 mM) and hyperglycemic (15 mM) exposures, focusing on hepatocellular mitochondrial function. HepG2 and HUH7 cell lines were exposed to CLEC and glucose for 24 weeks, mimicking a chronic heavy metal exposure paradigm seen in normal and type II diabetic individuals. We observe that CLEC exposures significantly affect the long-term health of mitochondria, including decreased mitochondrial mass, increased superoxide production, and loss of mitochondrial membrane potential (MMP) in a CLEC and glucose-dependent manner. Furthermore, the Seahorse MitoStress assay revealed CLEC induced significant chronic oxidative stress. In particular, CLEC cells showed altered levels of basal and non-mitochondrial respiration, causing dysregulation in mitochondrial oxygen consumption rates (OCRs). Lastly, we identified significant impacts of CLEC and glucose exposures on the mitochondrial dynamics (fission/fusion) of the CLEC cells, which showed enhanced mitochondrial fragmentation and turnover rates. We also identified novel cell compensatory mechanisms that may mask the true extent of chronic Cd exposure induced damage in liver cells. CLEC and glucose work additively to damage hepatocellular mitochondrial function. New approach methodologies (NAMs), such as the current vitro toxicology study, establish the insidious effects of chronic heavy metal pollutant exposures on human hepatocellular function.

## I. Introduction

Sustained heavy metal exposures are linked to a variety of human health morbidities. These include chronic diseases such as type II diabetes mellitus (T2DM), obesity, metabolic syndrome (MetS), cardiovascular disease, kidney diseases, and cancers. Humans are exposed to a wide variety of toxic heavy metals on a routine basis (Haidar et al., 2023). However, unlike organic xenobiotic pollutants that undergo enzymatic detoxification (e.g., through CYP450) (Reed & Backes, 2012), inorganic pollutants such as heavy metals tend to persist inside the body for a prolonged duration (i.e., decades) (Kumar & Gullapalli, 2024a; Wang et al., 2021), causing complex damage outcomes to cells and organs of the body (Haidar et al., 2023). Additionally, the chronic effects of heavy metal pollutants are ill-understood due to the diverse nature of the cellular damage inflicted by heavy metal accumulation within the body (Mitra et al., 2022). Heavy metals tend to preferentially accumulate in various organs of the body (e.g., cadmium (Cd) accumulation in the kidney and liver (Genchi et al., 2020) and lead (Pb) in the developing brain (Sanders et al., 2009)).

Heavy metal exposures are implicated in chronic metabolic diseases such as type II diabetes and obesity (Haidar et al., 2023). The incidence of human metabolic diseases is rapidly increasing across the globe for ill-understood reasons (Teng et al., 2023). Highly toxic metals, such as Cd, are implicated in the onset, progression, and severity of T2DM commonly seen in humans (Buha et al., 2020). Additional heavy metals such as lead (Pb) (Wang et al., 2018), mercury (Hg), arsenic (As), copper (Cu), and iron (Fe) also have a strong association with metabolic disorders such as type II diabetes, obesity, and metabolic syndromes (Haidar et al., 2023). T2DM in particular, is rising at an alarming rate worldwide, both in developed and developing countries such as India (Khan et al., 2020). The total number of T2DM patients in the United States (US) is estimated to be 38.4 million (around 11.6% of the population), and the number of pre-diabetics is estimated to be ∼ 97.6 million (around 38% of the population) (*National Diabetes Statistics Report*, 2024). Etiological factors driving T2DM include nutritional consumption imbalances (i.e., increased processed food consumption), sedentary lifestyles (i.e., lack of exercise), and inherited genetic factors, which have been well-studied (Ardisson Korat et al., 2014). And yet, the precise role of environmental pollutant exposures (e.g., heavy metals and chemicals) as drivers of anthropogenic metabolic disease is ill-understood, especially in the context of chronic heavy metal exposures.

The human morbidity and mortality associated with chronic metabolic dysfunction is mainly due to the accumulative damage seen in organs such as the liver (Vargas et al., 2024), kidneys (Bansal & Chonchol, 2025), and vascular systems (Swarup et al., 2025). In the liver, metabolic (dysfunction)-associated steatotic liver disease (MASLD; previously labeled non-alcoholic fatty liver disease - NAFLD) has become one of the most prevalent chronic liver diseases (Girish & John, 2025). MASLD is strongly associated with T2DM (up to 75% of type II diabetics have MASLD) (Dai et al., 2017), and it can further progress to metabolic steatohepatitis (MASH), fibrosis, cirrhosis, or hepatocellular carcinomas. Similar to T2DM, the etiopathogenesis of MASLD/MASH is multifactorial, though the precise molecular and cellular factors remain elusive (Miao et al., 2024; Younossi et al., 2025). There is increasing evidence that environmental exposures (such as heavy metal exposures) may play a key role in the increasing rates of MASLD/MASH observed globally (Kumar et al., 2025; Li et al., 2023).

Functionally, the liver is a key organ of human xenobiotic metabolism, detoxification, and systemic metabolic homeostasis (Kalra et al., 2025). Unsurprisingly, human livers are susceptible to hepatotoxicity due to a multitude of environmental toxin exposures, including heavy metals and chemical pollutants (Haidar et al., 2023). While the mechanisms underpinning hepatotoxicity of specific drugs are relatively well-understood (e.g., acetaminophen; Ramachandran & Jaeschke, 2017), the general hepatotoxic mechanisms of environmental pollutant exposures (e.g., heavy metals) remains poorly understood, particularly in the context of metabolic diseases such as T2DM.

Previous work from our lab established a novel *in vitro* model of heavy metal exposure effects on hepatotoxicity, which we termed as a “CLEC” model. Specifically, this *in vitro* model assessed the modulatory effects of chronic, low-dose exposures of cadmium (hence CLEC) on hepatocellular function, focusing on the insulin signaling pathway. We modeled the effects of pathophysiologically relevant levels (200 nM and 1 μM CdCl_2_) of Cd exposures on the insulin signaling pathway, focusing on the insulin-PI3K-Akt-mTOR signaling axis (Kumar & Gullapalli, 2024a). Dysfunction in this key cell signaling pathway underpins a variety of human metabolic dysfunctional disorders, including T2DM, obesity, MASLD, MetS, and even hepatocellular carcinomas (HCC). A key innovation of this model of chronic heavy metal stress was the assessment of the long-term effects of Cd with superimposed glucose levels, modeling the normo- (5.6 mM) and hyperglycemic (15 mM) states mimicking normoglycemic and T2DM conditions, respectively. A temporal dosing period of 24-week exposure to Cd was chosen to model the chronic exposure conditions (> 3 months) as per standard toxicology textbook definitions (Kumar & Gullapalli, 2024a, 2024b).

We utilize Cd to model chronic heavy metal stress exposures on liver cells to delineate specific molecular mechanisms of chronic heavy metal hepatotoxicity. Cd is a ubiquitous environmental heavy metal pollutant with no known physiological function in humans and is a pollutant of major global concern. Chronic Cd accumulation has a long biological half-life of 9–35 years in the human body (Kumar & Gullapalli, 2024a; Wang et al., 2021). More than 50% of Cd entering the human body bioaccumulates in the kidney or liver (Genchi et al., 2020). Due to long-term storage potential of Cd (and other heavy metals) in the liver (Alissa & Ferns, 2011; Genchi et al., 2020), understanding the modulatory impacts of CLEC and diabetes on chronic liver stress is of urgent concern. More broadly, there is a pressing need to understand the chronic impacts of heavy metal exposures such as Cd on liver pathophysiology, as a potential cause and modifier of human metabolic diseases such as T2DM and MASLD (Moroni-Gonzalez et al., 2023).

Pre-clinical testing for hepatotoxicity assessment is a prerequisite for human drug testing trials. Due to ethical issues and the “non-representativeness” of animal testing, there is an increasing push towards the use of human-relevant, pre-clinical models of drug development (e.g., new approach methodologies (NAMs), liver organoid models, liver-on-a-chip, 3-D bioprinting, stem cells, etc.) (Nair & Weiskirchen, 2024; Schmeisser et al., 2023). In the current study, we examine CLEC effects on hepatotoxicity, focusing on mitochondrial dysfunction specifically in one such NAM model. Our previous study identified elevated reactive oxygen species (ROS) production in CLEC and glucose-modulated cells (Kumar & Gullapalli, 2024a, 2024b). As a primary source of ROS species in cells, mitochondria are susceptible to damage due to divalent heavy metals (such as Cd) (Korotkov, 2023) and/or T2DM (Zhang et al., 2023). However, the precise mechanisms of chronic, sustained heavy metal stress on mitochondrial function have not been studied thus far, which is the major focus of this study. In addition to the CLEC-induced mitochondrial damage and hepatotoxicity, the current study also aims to shed light on the modulatory effects of glucose co-exposures on mitochondrial hepatotoxicity outcomes. In most therapeutic drug/xenobiotic studies in humans, a one-to-one mechanism (drug/chemical → hepatocellular damage) is used (Wang et al., 2025), with scant attention to modulatory real-world factors such as co-existing T2DM (e.g., do T2DM patients have a higher incidence of drug-induced liver injury reactions?) (Li et al., 2019). The current study aims to understand modulatory effects of chronic heavy metal exposures and/or T2DM on hepatic mitochondrial damage, leveraging our novel CLEC *invitro* NAM model to understand the role of divalent chronic heavy metal stress on hepatic metabolic dysfunction.

## II. Materials and Methods

### a. Chemicals and reagents

The main reagents utilized in this study for various assays are listed as follows: DMEM (Dulbecco’s Modified Eagle’s Medium) (Sigma # D6046), phosphate buffered saline (PBS; cat # P3813), D (+) Glucose solution (cat # RNBJ83621), and cadmium chloride (CdCl_2_; cat # 202908) were purchased from Sigma Aldrich (St. Louis, MO, US). MitoSox Red (cat# M36008), and Mitotracker green (cat # M7515), and Alamar Blue™ (cat #DAL1100), were purchased from Thermo Fisher Scientific, Waltham, MA. Rat tail collagen (cat # 5056) was purchased from Advanced Biomatrix (Carlsbad, CA), and RIPA lysis and extraction buffer (cat # 786–489) were purchased from G Bioscience (Saint Louis – US). Tetramethyl rhodamine ethyl ester (TMRE) (cat # 21426) and D-galactose (cat # 20890) were purchased from Cayman chemical (Ann Arbor, MI, US). All the reagents were used in their purchased form without further purification or modifications.

### **b.** Cell culture conditions and CLEC model development

HepG2 and HUH7 cell line models were purchased from American Type Culture Collection (ATCC; Manassas, VA). HepG2 cells were cultured in T25 and T75 tissue culture flasks coated with rat tail collagen (50 μg/ml). HUH7 cell lines were cultured in similar flasks without any modification. Chronic, low-dose exposures of Cd (CLEC) models used in the study were developed by exposing the HepG2 and HUH7 cell lines to cadmium chloride (CdCl_2_) in the following manner: untreated (0 nM), and 1 μM CdCl_2_ passaged serially for 24 weeks. Additionally, the Cd dosing concentrations (0 nM, and 1 μM) were used in conjunction with either normoglycemic (5.6 mM) or hyperglycemic (15 mM) cell culture conditions to mimic physiologically normal blood glucose levels (<100 mg/dL or ∼5.4% Hb1Ac) and uncontrolled diabetic states of humans (275 mg/dL or ∼11 % Hb1Ac) respectively (Fig 1). All cells were passaged 7-10 days, and the media was changed every 2–3 days serially for 24 weeks prior to making stocks. The cells were retrieved for the individual experiments conducted in the study as needed. All of the cell lines (under various CLEC exposure conditions) were cultured in a humidified incubator (Thermo Fisher Scientific, Waltham, MA) maintained at 5 % CO2 and 37 ◦C temperature for the duration of the exposure.

**Fig. 1.**
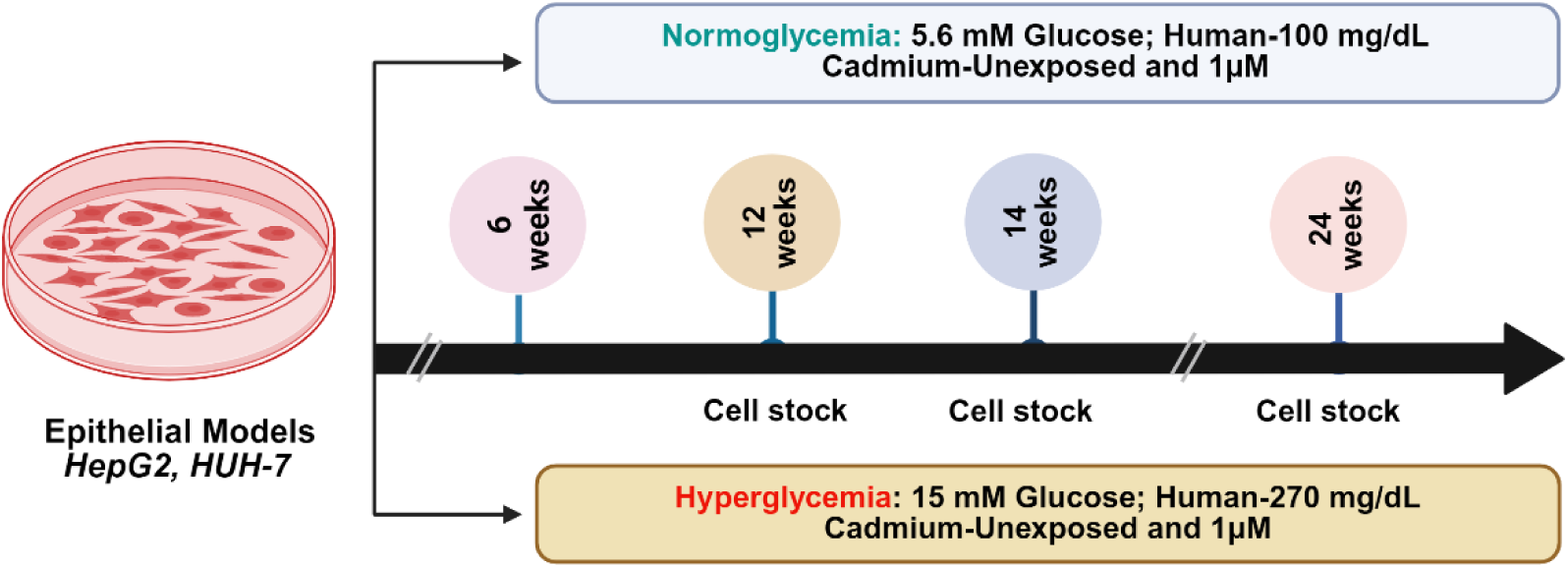
The CLEC exposure paradigm. Epithelial liver cell lines (A) HepG2 and (B) HUH7 were exposed to varying glycemic conditions (normoglycemia 5.6 mM and hyperglycemia 15 mM) with/without added cadmium chloride (CdCl_2_) for a total duration of 24 weeks. At the 24-week time point, cell stocks were prepared and preserved in liquid nitrogen. A new stock vial is revived one week prior to various experiments.

### **c.** CLEC cell viability evaluation using dose-response curve analysis

HepG2 and HUH7 cells were seeded at a density of 7,000 cells per well in 96-well plates and allowed to adhere overnight in their respective glycemic conditions (normoglycemia 5.5 mM and hyperglycemia 15 mM), supplemented with 10% fetal bovine serum (FBS). Before seeding the HepG2 cells, the 96-well plates were pre-coated with rat tail collagen and incubated at room temperature for 2 hours, followed by three washes with phosphate-buffered saline (PBS). The next day, cells were treated with progressively increasing concentrations of Sorafenib (0, 0.5, 1, 5, 10, 25, 50, and 100 µM) in serum-free media under either normoglycemic (5.5 mM) or hyperglycemic (15 mM) conditions. After 12 hours, 10 % Alamar Blue™ of total volume was added to each well, and absorbance readings were taken after 4-5 hours at 570 and 600 nm wavelengths using the Bio-Tek Synergy Neo2 Plate reader (Bio-Tek Instruments, Inc, Winooski, VT, USA). Dye reduction absorbance was measured to calculate the overall cell viability in each well according to the manufacturer’s instructions to determine the final EC5, EC10 and EC50 values. Cell viability data are shown as mean +/-standard deviation of at least two temporally independent experiments with five individual replicates of each test condition in the 96-well plate. A four-parameter, log-logistic model was used to compute cell viability of the cells using the “drc” package from R. The non-linear regression equation used for the model fit is given as follows

Where,

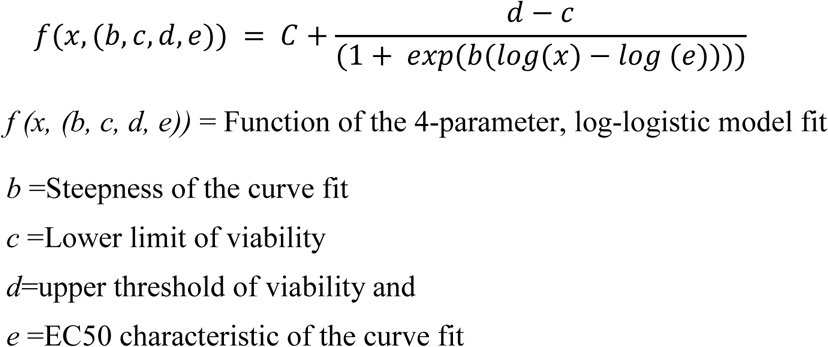

### **d.** CLEC cell viability assessment using galactose selective media conditions (Glu/Gal assay)

To determine the CLEC cell dependence on oxidative phosphorylation (as opposed to mainly glycolysis), we performed further cell viability analyses (as above) with galactose media substitution. This well-established assay is commonly referred to as the Glu/Gal assay and is routinely used in evaluation of invitro drug toxicity (Swiss & Will, 2011). CLEC cells were seeded at 7,000 cells/well as before and allowed to adhere overnight under regular glycemic conditions (normoglycemia 5.5 mM or hyperglycemia 15 mM), supplemented with 10% fetal bovine serum (FBS). After overnight incubation, the glucose containing culture medium was replaced with serum- and glucose-free Dulbecco’s Modified Eagle Medium (DMEM) containing 10 mM galactose. Cells were preconditioned in the galactose-DMEM medium for 2 hours to facilitate metabolic adaptation. Following the preconditioning phase, cells were then exposed to increasing concentrations of Sorafenib (0, 0.1, 0.25, 0.5, 1, 5, and 10 µM doses) in the serum- and glucose-free DMEM supplemented with 10 mM galactose. After a 10-hour incubation period, 10% Alamar Blue™ solution was added to each well, and absorbance readings were collected after 6-7 hours (total treatment period – 16-17 hours) at 570 and 600 nm wavelengths using a Bio-Tek Synergy Neo2 Plate Reader (Bio-Tek Instruments, Inc., Winooski, VT, USA). Cell viability and fit were calculated as before using a 4-parameter log-logistic model.

### **e.** Superoxide radical measurements

Mitochondrial superoxide generation was evaluated using fluorescent probe MitoSox Red (Thermo Fisher Scientific, cat # M36008). Oxidation of the MitoSox probe yields intermediate, probe-derived radicals that are successively oxidized to generate the corresponding fluorescent products. CLEC cells were plated at 5,000 cells per well into 96-well plates in 200 µL of growth medium with respective to their glycemic and CdCl_2_ concentrations. Next day, the media was replaced, and cells were stained with prewarmed media with 200 nM of MitoSox Red for 30 min in dark conditions at 37°C. Subsequently, cells were washed with PBS three times and stained with the Hoechst 33452 (ThermoFisher Scientific, cat # H1399) for 10 min. After incubation, the total superoxide radical induced fluorescence intensity was measured using the high throughput Cellomics CX7 instrument (ThermoFisher Scientific, Waltham, MA). Fluorescence intensity of 1000 cells were measured in each well and normalized on a 0 to 1 scale. Three, temporally independent, replicates were performed with a total of five replicate wells for each exposure condition (n=15) to measure the superoxide radical production in each CLEC cell condition.

### **f.** Measuring mitochondrial mass and morphological assessments

To evaluate differences in the mitochondrial mass and morphology due to the CLEC exposures, fluorescence imaging-based measurement of the mitochondria was performed on a high-content screening (HCS) platform, Cellomics CX7 (ThermoFisher Scientific, Waltham, MA). Mitochondrial mass was measured on the HCS platform by incubating the CLEC cells with 200 nM Mito-Tracker Green (Thermo Fisher Scientific, cat # M7515) in fresh growth medium (DMEM) prewarmed at 37 °C for 30 min. 5,000 cells of the different CLEC exposure conditions were seeded in a 96-well plate with specific DMEM media conditions and CdCl_2_ overnight. After overnight incubation, cell media was changed and washed once with PBS. Subsequently, cells were treated with 200 nM Mito-Tracker Green dye in prewarmed DMEM and incubated at 37 °C for 30 mins under dark conditions. After the incubation period, cells were washed three times with PBS, and the total fluorescence intensity was measured using the high-throughput Cellomics CX7 (ThermoFisher Scientific, Waltham, MA) platform. Fluorescence intensity from 1000 cells/well was measured and normalized within CLEC condition. Evaluation was done at three separate time points with a total of five replicates (total n=15) for each CLEC exposure conditions.

Confocal microscopy (Zeiss LSM 900, ZEN 3.7, Zeiss Optical, Jena) was performed next to evaluate the CLEC mitochondrial morphological differences in detail. 50,000 cells were seeded on four well chamber slides (Thermo Scientific Nunc Lab-Tek II Chamber Slide cat # 154526) at the respective glycemic (5.6 mM and 15 mM) and cadmium (0 and 1 µM) concentrations. CLEC cells were fixed with 4% paraformaldehyde for 10 minutes at room temperature. After blocking, cells were washed with PBS followed by blocking and permeabilization using 5% BSA + 0.03% Triton X-100 for 1 hour at room temperature. For nuclear staining DAPI (Thermo Scientific cat # D1306) was used. The cells were probed with the TOMM20 antibody delineate the mitochondrial morphology by incubating the fixed cells at 4°C overnight. The cells were washed next day with PBS and incubated with fluorescence-tagged secondary antibodies for 2 hours with constant shaking in the dark at room temperature prior to imaging. Images were captured with the Zeiss confocal microscope using a 63x oil immersion objective.

### **g.** Mitochondrial Membrane Potential assessment

The baseline differences in the CLEC cell mitochondrial membrane potential (MMP) were evaluated by measuring the uptake of the Tetramethyl rhodamine ethyl ester (TMRE) dye (Cayman Chemical, cat # 21426). TMRE is a cell-permeant, cationic, red-orange, fluorescent dye readily sequestered by active mitochondria depending on the mitochondrial membrane potential gradient. CLEC cells were seeded at a density of 5000 cells/well in a 96-well plate and allowed to grow overnight for baseline MMP assessment. The cells were incubated next day with 100 nM TMRE (final concentration) in prewarmed DMEM (without phenol red) for 30 min at 37°C in a CO_2_ incubator under dark conditions. After incubation, cells were washed three times with PBS and then stained with the Hoechst 33452 nuclear stain for a further 10 min. TMRE fluorescence intensity from 1000 cells/well was measured by acquiring images on the high-throughput Cellomics CX7 platform (ThermoFisher Scientific, Waltham, MA) at an excitation wavelength of 545 nm. The fluorescence emission was collected using the Cellomics instrument filter set (549-15_BGS-RS (590 nm) where 549-15 represents the excitation, BGS-dichroic mirror, and RS is the emission filter). The MMP assay was performed at three separate time points with a total of five replicate wells for each time point across the different CLEC exposure conditions (n=15; 75,000 cells in total). Quantitative differences in the intensity of the TMRE uptake between the individual CLEC conditions was calculated to determine baseline MMP differences.

### **h.** Metabolic Assay Measurements (Oxygen Consumption Rate and Extra-Cellular Acidification Rate)

The real time monitoring of the mitochondrial respiratory status of the CLEC cells was measured using the Seahorse XFe24 Analyzer platform (Agilent Bioscience, Billerica, MA, USA). Specifically, the Seahorse XF Cell Mito Stress Test Kit (catalog number #103015-100, Agilent Technologies, Santa Clara, CA, USA) was used to determine the differences in the oxygen consumption rates (OCR) and extra-cellular acidification rates of CLEC cells at baseline according to manufacturer’s instructions. Cells were seeded at density of 4 × 10^5^ cells/well (after optimization) in 500 μL of DMEM under standard glucose and Cd incubation conditions. After 24 hours of seeding, the growth medium was replaced with 500 μL/well of Seahorse XF DMEM assay medium containing 1 mM pyruvate, 2 mM glutamine, and the specific glycemic conditions (normoglycemia 5.5 mM and hyperglycemia 15 mM which was maintained consistently throughout the experiment). The Seahorse plates were incubated next day in a 37°C non-CO2 incubator for 1 h, before starting the experimental procedure.

The sensor cartridge was hydrated using Seahorse XF calibrant and incubated at 37°C in a non-CO_2_ incubator for overnight. For the Mito stress assay, mitochondrial complex inhibitors and uncouplers, oligomycin, FCCP, rotenone and antimycin A (part of the kit) were made in the prewarmed seahorse media and sequentially loaded into injector ports A, B and C of sensor cartridge, at working concentrations of 1 μM, 1 μM, and 0.5 μM, respectively. A nuclear stain (Hoechst 33452) was loaded in the D port to count the number of cells and normalize the data. The OCR and ECAR values of CLEC cells were measured under basal conditions. Additionally, derivative non-mitochondrial respiration, maximal respiration, proton leak, ATP respiration, respiratory capacity, coupling efficiency values were measured in the assay according to the manufacturer guidelines. Three temporally independent replicates of the Mito stress assay were performed. The OCR and ECAR data were analyzed using Wave software v.2.6 (Agilent) in the online and offline mode. The resulting Seahorse assay data was plotted using the GraphPad Prism software (GraphPad prism 9, Boston, MA, US).

### **i.** RT-PCR analysis of anti-oxidant pathway genes

Relative gene expression levels of select mitochondrial and cytoplasmic antioxidant genes were determined using a quantitative real-time reverse transcriptase polymerase chain reaction (qRT-PCR) method. For each of the CLEC conditions, the supernatant media was removed, and the cells were washed with cold 1X PBS, followed by the addition of 800μl of Trizol IBI isolate (cat # IB47600, IBI Scientific, Iowa) directly onto the cells in T-25 flasks. The cell supernatant was pipetted and transferred into a 1.5ml Eppendorf tube followed by addition of 200μl chloroform. The cell solution was subsequently spun down for 15mins at 14,000rpm. After centrifugation, the top clear/yellowish layer was isolated for RNA retrieval, followed by transfer to a new Eppendorf tube and addition of 500μl isopropanol. The mixture was incubated for 15 mins at room temperature followed by a second step of centrifugation at 14,000 rpm for an additional 15 mins. Finally, 75% EtOH was added to the pellet and centrifuged at 14,000 rpm for 10 mins. The ethanol was aspirated, and the pellet was resuspended in diethyl pyrocarbonate (DEPC) water (Cat# AM9915G, Thermo Fisher Scientific, Waltham, MA). The purified RNA was quantified using a Nanodrop instrument (Thermo Fisher Scientific, Waltham, MA). One μg of purified RNA was reverse transcribed using Thermo Scientific

Revert Aid First Strand cDNA synthesis kit (Cat# K1621 Thermo Fisher Scientific, Waltham, MA). Applied Biosystems Fast SYBR Green Master Mix (cat# 4385612 Thermo Fisher Scientific, Vilnius, Lithuania) was used to quantify the mRNA gene expression. Gene expression was normalized to ß-Actin gene expression used as an internal control gene standard. The qPCR primers were ordered from Integrative DNA technologies (IDT, San Diego, CA, US). The relative gene expression was quantified using the 2^-△△CT^ method. The list of genes and primers sequences used for the qRT-PCR analysis are listed in table 1.

**Table 1.**
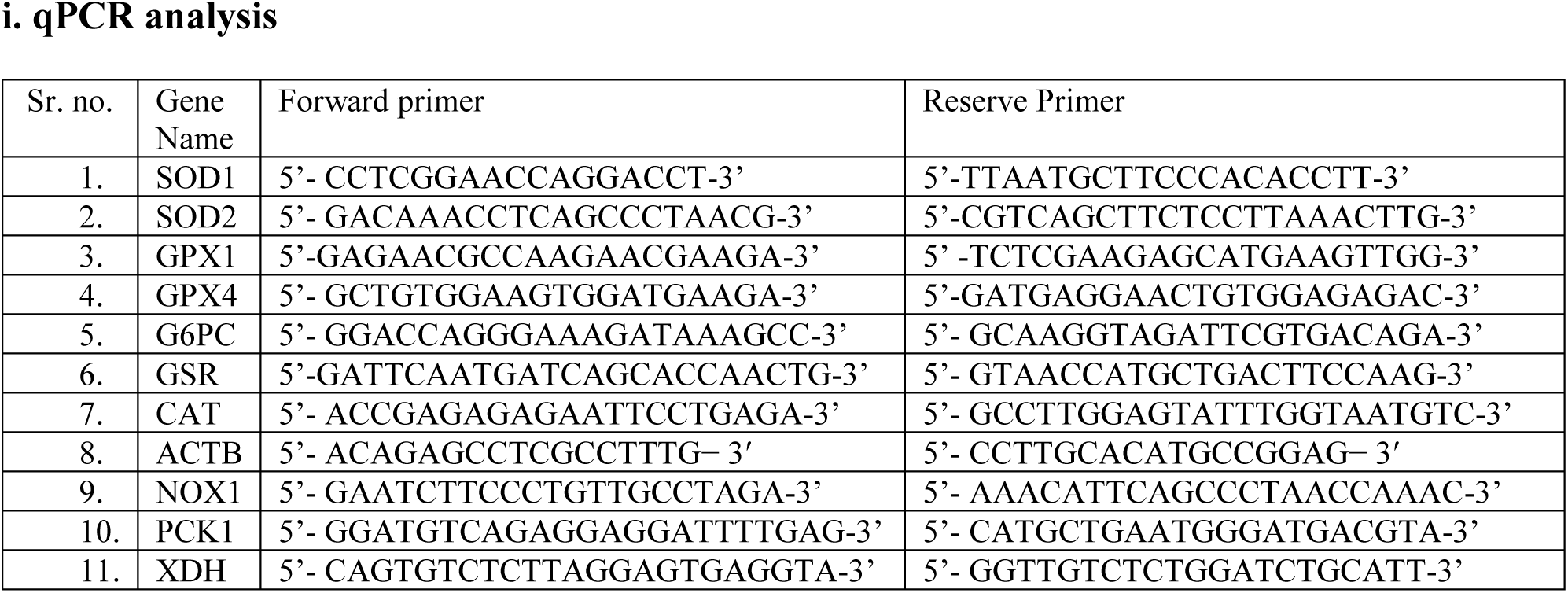
the list of gene and primer sequence used in the study for the qRT-PCR analysis.

### **j.** Western blot

Cell proteins were analyzed using standard Western blot protocols. In brief, CLEC exposure cells were seeded on 100mm Petri dishes. Cells were washed with PBS and lysed in RIPA buffer containing protease and phosphatase inhibitors (cat # A32959, Thermo Fisher Scientific, Carlsbad, CA). Subsequently, cell extracts were prepared by trypsinization followed by centrifugation at 14000 rpm (at 4℃) for 15 minutes. Cell extract protein concentrations were quantified using the Pierce™ protein assay reagent (cat # 23227 Thermo Fisher Scientific, Carlsbad, CA). Proteins (30μg) were separated on a 4% −12% Bis-Tris-SDS gel (cat # 4561094, Bio-Rad, Hercules, CA, USA) and transferred to a nitrocellulose membrane, 0.45μm (cat #162–0115, Bio-Rad Laboratories Inc. Hercules, CA). The membrane was blocked using a skimmed milk suspension for one hour and incubated with the primary antibodies (1:1000 dilution) overnight at 4◦C. The list of primary and secondary antibodies used in the study are listed in table 2. Western blots were washed with Tris-buffered saline with 0.1% Tween 20 (TBST). The membranes were incubated further with the secondary antibody (1:5000) for 1 hour at room temperature. Finally, the Western blots were developed using the ECL chemiluminescence detection reagent (cat # 34076, Thermo Fisher Scientific, Carlsbad, CA). Western blots were imaged using on a ChemiDoc imaging station (Bio-Rad, Hercules, CA). Bio-Rad Image lab software was used to quantify the density of the Western blot bands for quantitative ratiometric assessments using a beta-actin control. Western blots for each exposure condition were performed at baseline extracted from independent samples in triplicate (n=3) to confirm the protein expression changes reported in the CLEC cells.

**Table 2.**
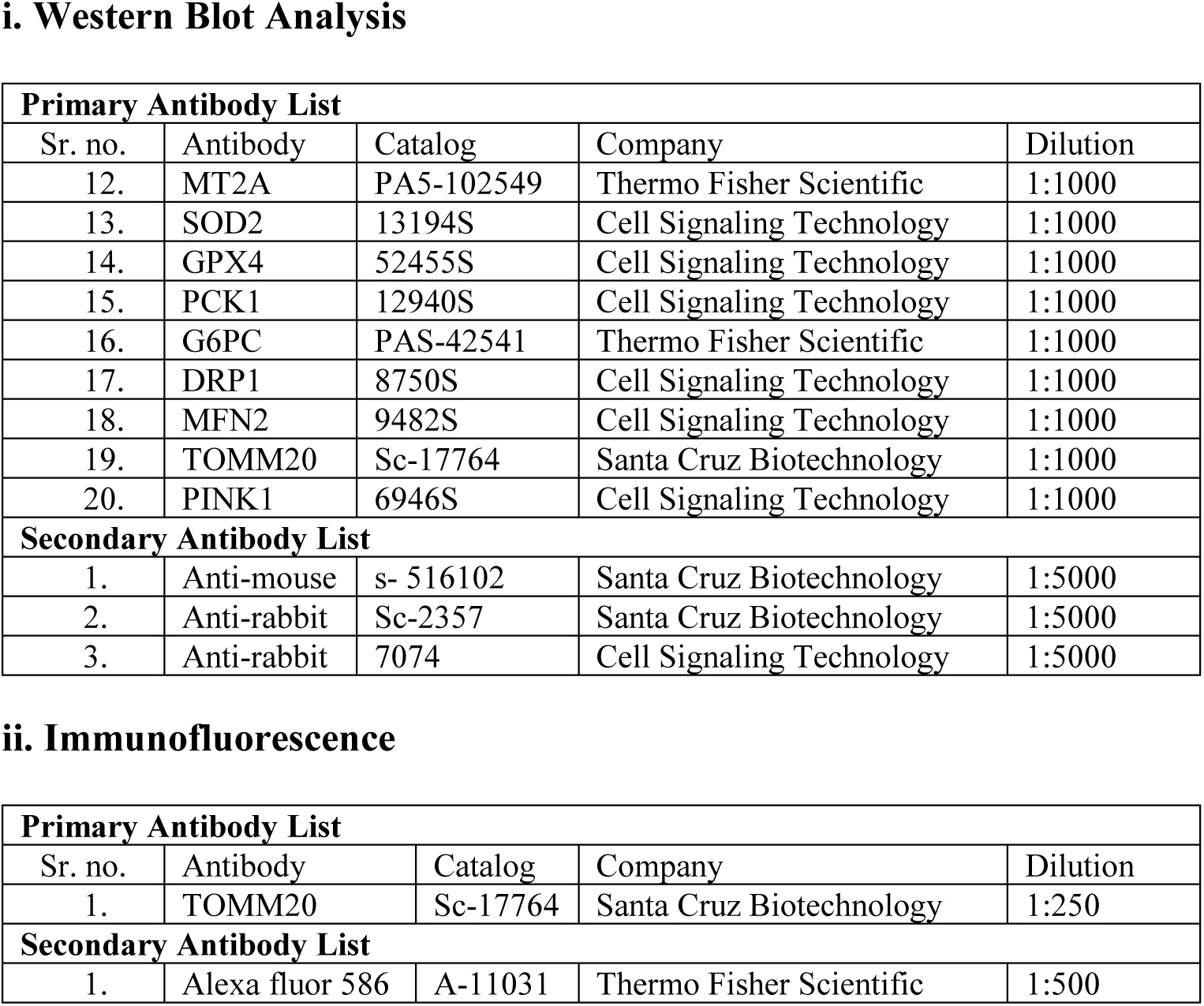
List of primary and secondary antibodies used for Western blot analysis and immunofluorescence staining.

i. Western Blot Analysis
| Primary Antibody List |  |  |  |  |
| --- | --- | --- | --- | --- |
| Sr. no. | Antibody | Catalog | Company | Dilution |
| 12. | MT2A | PA5-102549 | Thermo Fisher Scientific | 1:1000 |
| 13. | SOD2 | 13194S | Cell Signaling Technology | 1:1000 |
| 14. | GPX4 | 52455S | Cell Signaling Technology | 1:1000 |
| 15. | PCK1 | 12940S | Cell Signaling Technology | 1:1000 |
| 16. | G6PC | PAS-42541 | Thermo Fisher Scientific | 1:1000 |
| 17. | DRP1 | 8750S | Cell Signaling Technology | 1:1000 |
| 18. | MFN2 | 9482S | Cell Signaling Technology | 1:1000 |
| 19. | TOMM20 | Sc-17764 | Santa Cruz Biotechnology | 1:1000 |
| 20. | PINK1 | 6946S | Cell Signaling Technology | 1:1000 |
| Secondary Antibody List |  |  |  |  |
| 1. | Anti-mouse | s- 516102 | Santa Cruz Biotechnology | 1:5000 |
| 2. | Anti-rabbit | Sc-2357 | Santa Cruz Biotechnology | 1:5000 |
| 3. | Anti-rabbit | 7074 | Cell Signaling Technology | 1:5000 |

ii. Immunofluorescence
| Primary Antibody List |  |  |  |  |
| --- | --- | --- | --- | --- |
| Sr. no. | Antibody | Catalog | Company | Dilution |
| 1. | TOMM20 | Sc-17764 | Santa Cruz Biotechnology | 1:250 |
| Secondary Antibody List |  |  |  |  |
| 1. | Alexa fluor 586 | A-11031 | Thermo Fisher Scientific | 1:500 |

### h. Statistical assessment

All data in this study is represented as mean ± standard deviation (S.D) for different quantitative assessment values. Statistical analysis to determine differences in the CLEC exposures were performed using a two-way ANOVA analysis design with i. Cd, ii. Glucose and iii. Cd x Glucose interactions as the key factors assessed. Subsequently, we performed a post-hoc Tukey’s multiple comparison test to identify statistically significant mean differences between groups. A value of p < 0.05 is considered significant for the statistical test difference. Values of statistical significance are represented as follows - p <0.05 (*), p<0.01(**), p<0.001(***), p<0.0001 (****) in the graphs. A minimum of three independent experimental replicates were performed for all assays with multiple technical replicates in each assay as appropriate and as described within the methods section describing each assay (see above).

## III. Results

### a. Measuring baseline cell viability differences in CLEC cells using mitotoxic Sorafenib

To determine effects of Cd and glucose exposures on hepatocellular mitochondrial function, we performed a dose-response curve (DRC) analysis using the mitotoxic drug, Sorafenib. Dose dependent mitotoxicity of Sorafenib (and multi-kinase inhibition) is well established in the literature. Study HepG2 and HUH7 CLEC exposures are labelled as follows - NG-UN (5.6 mM glucose, no Cd), NG-1µM (5.6 mM glucose, 1µM Cd), HG-UN (15 mM glucose and no Cd), and HG-1µM (15 mM glucose and 1µM Cd).

CLEC cells were treated with increasing concentrations of Sorafenib (0, 0.5, 1, 5, 10, 25, 50, and 100 µM; see methods). All CLEC cells demonstrated an anticipated dose-dependent reduction in cell viability at the end of a 16-to-17-hour Sorafenib exposure period (Fig 2). Differences in the EC_50_ values establish the effects of CLEC and glucose exposures on the cyto- and mitotoxic outcomes of Sorafenib. The EC_50_ values due to Sorafenib toxicity are approximately similar in the different CLEC exposures (Fig 2). HepG2 cells showed EC_50_ values ranging from 4.67 µM to 2.85 µM while the HUH7 CLEC cells range from 2.78 µM to 1.62 µM (see supplementary table 1 for specific EC_50_ values). Overall, the HepG2 CLEC model is less susceptible to Sorafenib induced toxicity compared to the HUH7 CLEC cell line. However, the cytotoxicity EC50 patterns in the individual CLEC cell models (i.e., NG-UN vs NG-1 µM vs HG-UN vs HG-1 µM) demonstrates a mixed pattern (Fig 2).

**Fig. 2.**
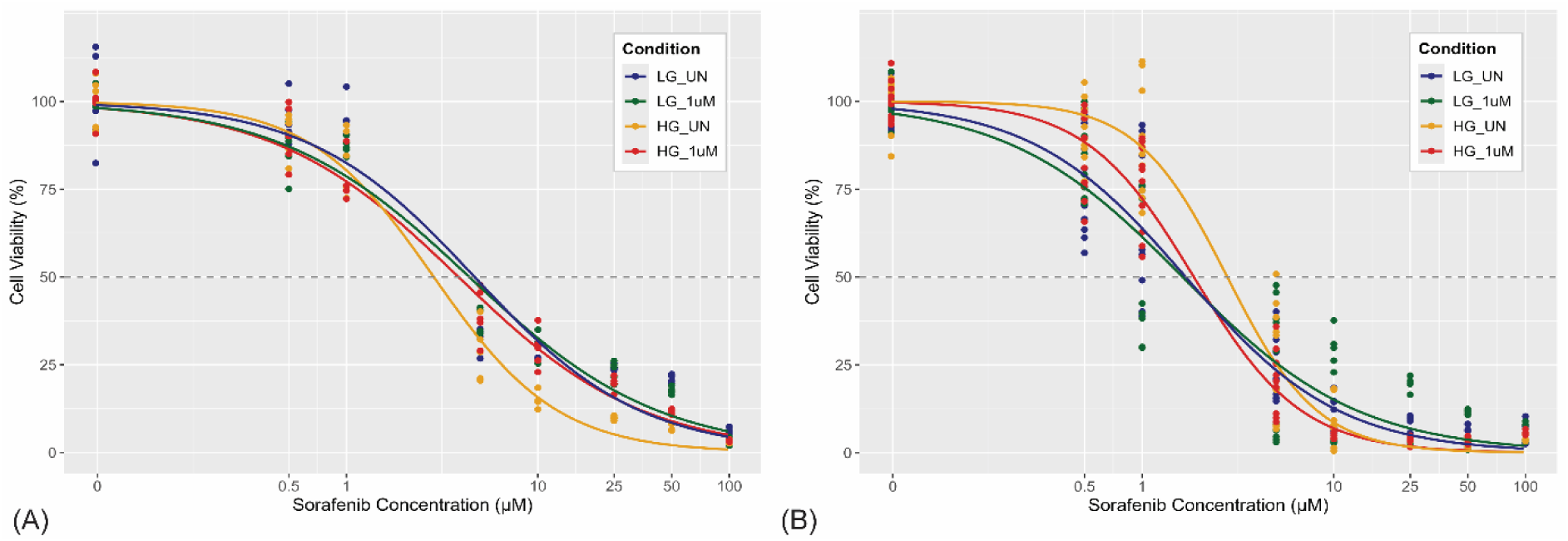
Dose-response curve (DRC) analysis of A. HepG2 and B. HUH7 CLEC cells using mitotoxic Sorafenib. CLEC and glucose conditioned cells were exposed to varying concentrations of Sorafenib (0, 0.5, 1, 5, 10, 25, 50, and 100 µM) under glycemic (5.6 mM or 15 mM) cell culture conditions to determine the lethal concentration (EC_50_) for a period of 16 hours. Different conditions are represented by the different colors. DRC shows minor differences in the EC_50_ values in the CLEC and glucose exposed cells in response to acute exposures of Sorafenib. These experiments were performed in duplicate (n=2) with five replicates in each plate and the data was fit using a four-parameter, log-logistic model using the R package, DRC.

The HepG2 cell normoglycemic (NG-UN) condition shows least susceptibility to Sorafenib compared to the hyperglycemic state (HG) as expected. In contrast, this trend is slightly reversed in the HUH7 CLEC exposure model. We postulate these minor differences (and lack of an overall observable trend) in the EC_50_ values are likely due to adaptive compensation occurring over the course of the long duration of the Cd and glucose exposures (24 weeks) and the intrinsic experimental variability associated with the dose-response curve assay.

### **b.** Glu/Gal assay unmasks mitochondrial dysfunction due to CLEC and glucose exposures

Glucose metabolism and adaptive compensation may mask the true mitochondrial toxicity (i.e., protective glycolysis due to Crabtree effect) over the course of a 24-week exposure. Consequently, the true impact of CLEC and glucose exposures on mitochondrial dysfunction may be masked with acute Sorafenib mitotoxic exposure. To understand the mitochondrial specific impacts of CLEC and hyperglycemia, we performed the Glu/Gal DRC assay. In brief, glucose culture media is substituted with galactose (10 mM; see methods) followed by Sorafenib dose-response curve assessment. This approach is the well-established *Glu-Gal assay* technique which imposes a state of forced oxidative phosphorylation on cells as galactose is preferentially metabolized by mitochondria without entering the glycolytic cycle. Under such galactose exposure conditions, CLEC cell models are forced to use the dysfunctional mitochondria for survival upon which, the Sorafenib toxicity is superimposed. We observe the cell viability is significantly reduced across all CLEC conditions under forced oxidative phosphorylation (Fig 3). Results are presented as EC_50_ values in each CLEC condition and the Glu/Gal ratios (EC_50_ values in Glucose media/EC_50_ value in galactose media; see supplementary table 1). The Glu/Gal ratios range from 5.1 to 8.99 in HepG2 CLEC cells and 11.7 to 40 in HUH7 cells indicating significant cell death with galactose media. More importantly, the Glu/Gal analysis unmasks the long-term effects of Cd and Glucose exposures on mitochondrial function in CLEC cells.

**Fig. 3.**
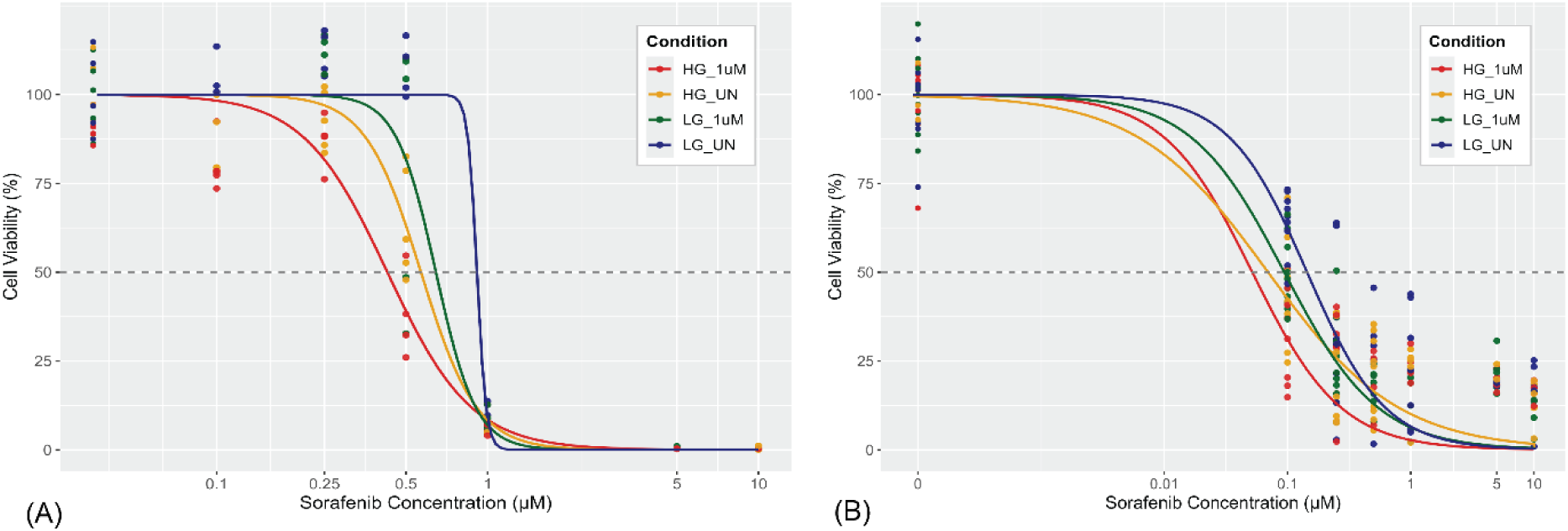
Glu/Gal Dose-response curve (DRC) analysis of A. HepG2 and B. HUH7 CLEC cells – CLEC cells are exposed to mitotoxic Sorafenib with galactose media replacement (Glu/Gal assay). All the conditions (NG-UN, NG-1µM, HG-UN, HG-1µM) are exposed to varying concentrations of Sorafenib (0, 0.1, 0.25, 0.5, 1, 5, and 10 µM) in serum and glucose free media with 10 mM galactose to determine the galactose modified EC_50_ values. The experiments were performed in duplicate (n=2) with the five replicates per condition and the DRC data was fit using a four-parameter, log-logistic model using the R package, DRC.

In the HepG2 CLEC cells, we observe the EC_50_ values are lowest in hyperglycemia-1µM Cd cells (EC_50_ - 0.43 µM) indicating significant mitochondrial dysfunction while the normoglycemic, untreated cells are most resistant to Sorafenib (EC_50_ – 0.91 µM). The EC_50_ values in the HepG2 CLEC cells follow the trend of NG-UN > NG-1µM > HG-UN > HG-1µM as expected. Similar to HepG2, In HUH7, the highest EC_50_ value is noted with normoglycemia-unexposed condition (EC50 value – 0.14 µM) while the lowest EC_50_ value is seen in the hyperglycemic state with and without Cd (EC50 – 0.05 and 0.07 µM, respectively). The HUH7 EC_50_ trend mirrors the HepG2 pattern as follows - NG-UN > NG-1µM > HG-UN > HG-1µM. We observe cell-line specific differences (i.e., HepG2 is resistant to cell death compared to HUH7; see supplemental table 1 for EC_50_ values). These results establish the HUH7 CLEC cell line demonstrates an enhanced sensitivity to Sorafenib compared to HepG2 CLEC cells.

More importantly, these findings establish the impacts of chronic low-dose cadmium (CLEC) and hyperglycemic exposures as significant, long-term modulator of mitochondrial dysfunction and mitotoxicity. The DRC results illustrate the critical importance of chronic exposures (i.e., Cd and Glucose) on mitochondrial dysfunction in response to hepatic mitotoxic drugs. Thus, variable mitochondrial function is a key consideration in individuals with a history of heavy metal exposures, type II diabetes and metabolic dysregulation.

### **c.** CLEC cell models show differential mitochondrial morphology and mass at baseline using qualitative and quantitative approaches

The cell viability assays reveal mitochondria specific dysfunction due to chronic Cd and glucose exposures. To further establish the effects of CLEC and glucose exposures on mitochondria, we qualitatively and quantitatively measured the mitochondrial morphology and mass in the CLEC cell models. Qualitative evaluation of the mitochondria was performed using TOMM20 antibody labeling. TOMM20 is a key constitutive protein of the mitochondrial transport complex located on the outer membrane of mitochondria. Mitochondrial mass was semi-quantitatively assessed using the fluorescence dye, Mitotracker Green (see methods).

Confocal imaging of the TOMM20 staining reveals mitochondria in untreated cells (NG-UN) show an elongated and branching pattern of mitochondrial tubular networks in both HepG2 and HUH7 cells. Such patterns indicate normal and healthy mitochondrial dynamics and turnover (Fig 4). In contrast, the CLEC and/or glucose exposed cells (NG-1 µM, HG-UN, and HG-1 µM exposures) demonstrate a shift toward increased fragmentation of the mitochondria with edematous (swollen) and punctate appearing mitochondria, suggestive of impaired mitochondrial morphology and dynamics (Fig 4). The fragmented mitochondrial morphology is readily apparent in the CLEC exposure cells. The edematous appearance (swelling) of the mitochondria is observed to be greater in the hyperglycemic conditions compared to normoglycemia. Mitochondrial morphology differences are seen prominently in the HUH7 cell line compared to HepG2 suggesting cell line specific differences. TOMM20 staining also shows an increased perinuclear accumulative pattern due to CLEC and/or glucose exposures. This is a novel observation in liver cells not reported previously to our knowledge. Perinuclear accumulation of mitochondria is usually seen in conditions associated with increased iron stress commonly seen in hematological disorders such as sideroblastic anemias (Sheftel et al., 2009) (see discussion).

**Fig. 4.**
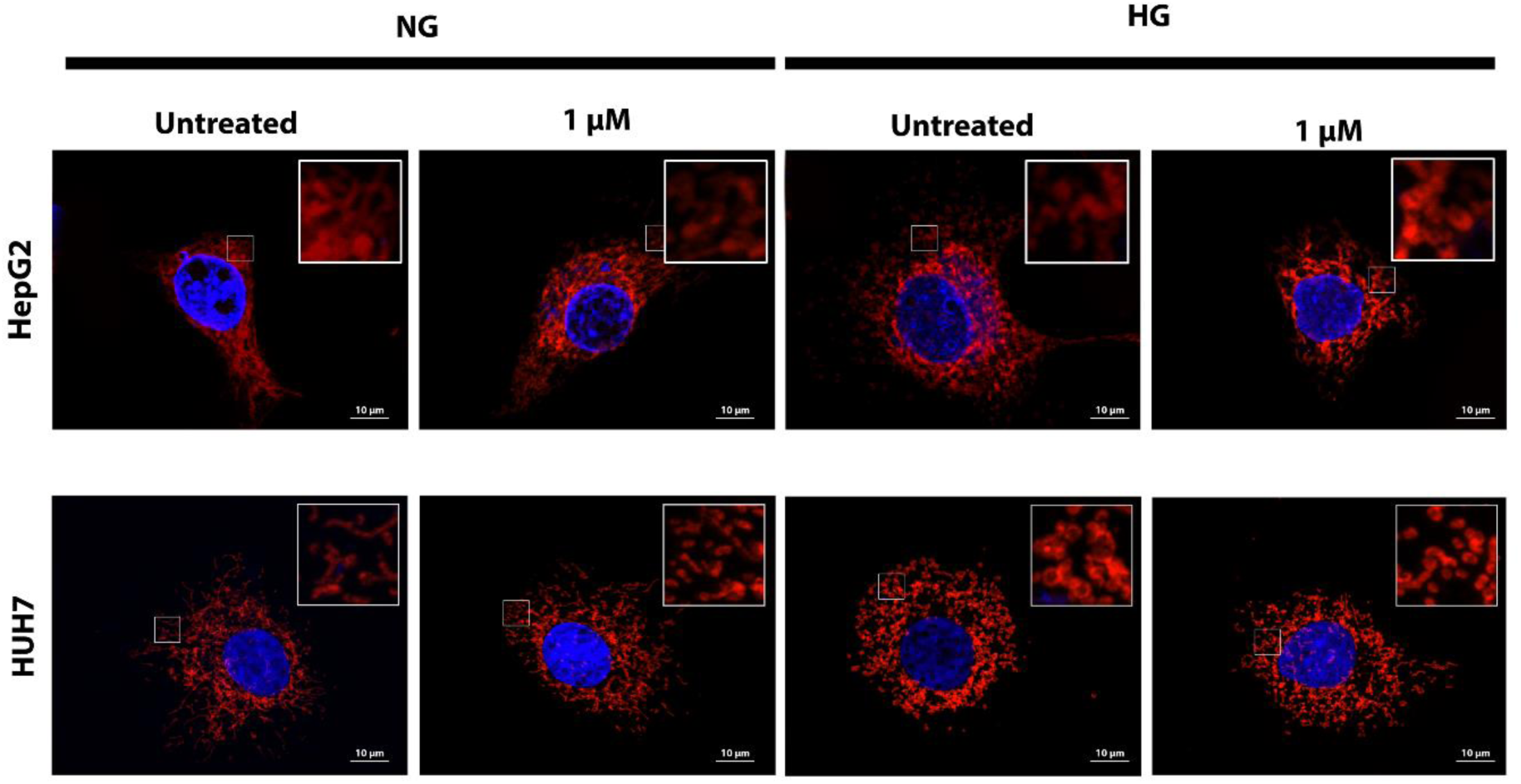
Representative confocal microscopy images of CLEC cell mitochondria – CLEC cells are stained with TOMM20 antibody (see methods). In both CLEC models, HepG2 (top row) and HUH7 (bottom row), untreated cells (no Cd and 5.6 mM glucose) show normal mitochondrial networks and filamentous morphologies (left). Adding Cd (1 µM) or hyperglycemia (15 mM) leads to altered morphology of the mitochondria. (Insets) Enlarged views of each CLEC condition highlight the various mitochondrial morphological alterations observed (edema, fragmentation and punctate appearance) seen in different CLEC cell conditions.

To robustly establish the qualitative mitochondrial staining patterns, we performed quantitative measurement of mitochondrial mass using the Cellomics high-content imaging system. As shown in figure 5, we observed a linear reduction in the total Mitotracker Green fluorescence intensity with Cd exposure, both in normoglycemia and hyperglycemia. This effect is observed consistently in both the CLEC cell models (HepG2 and HUH7). Additionally, under conditions of hyperglycemia alone (without Cd), a significant reduction in fluorescence intensity (up to 1.5-fold) is detected compared to normoglycemia. This effect is observed in both CLEC cell lines, suggesting Cd- and glucose-exposure dependent reduction mitochondrial mass in a chronic exposure setting. Two-way ANOVA analysis showed a statistically significant interaction between glucose and cadmium treatments on mitochondrial mass, emphasizing the toxic effects of combined CLEC and hyperglycemia on mitochondrial health (see supplementary table 1). In summary, CLEC exposures disrupt hepatocellular mitochondrial morphology and integrity, which is exacerbated in diabetic (hyperglycemic) environments.

**Fig. 5.**
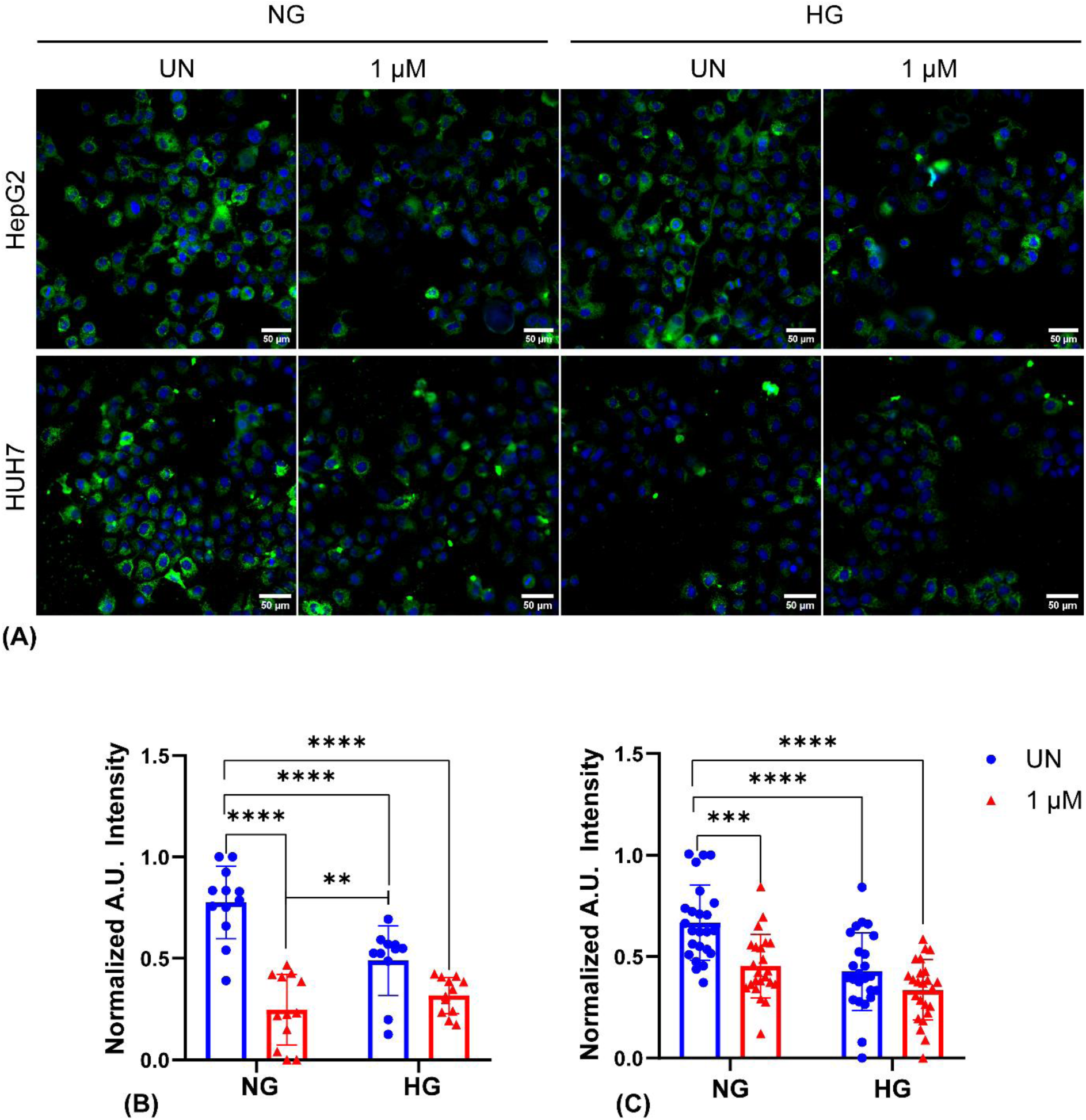
Quantitative measurement of mitochondrial mass in CLEC and glucose exposures – (A) CLEC cells were stained at baseline with 200 nM Mitotracker Green for 30 min. Both CLEC models (B) HepG2 and (C) HUH-7 show a significant decrease in the total mitochondrial mass in CLEC and hyperglycemic cell culture conditions. Hyperglycemia 1 µM Cd (HG-1 µM Cd) condition shows the least dye uptake indicating decreased mitochondrial mass. All the values are normalized using the max – min normalization method on a 0-1 scale. Data points on the bar plot represent a mean of three independent experiments with five well replicates in each plate (see methods). Statistical level of significance is described as follows - *p<0.05, **p<0.01, ***p<0.001, ****p<0.0001.

### **d.** CLEC exposures drive increased oxidative stress through enhanced superoxide radical production

Oxidative stress plays a key role in MASLD pathogenesis irrespective of the proximal drivers of the condition (Tauil et al., 2024). Acute exposures to Cd are well-known to induce oxidative stress through excess production of reactive oxygen species (ROS), including superoxide anions (O₂•⁻), hydroxyl radicals (•OH), and hydrogen peroxide (H₂O₂) (Yang et al., 2022; Zhang et al., 2020). However, CLEC exposure effects on mitochondria induced oxidative stress are lacking currently which was measured in this assay. At baseline, we measure highest superoxide levels in the HG-1 µM Cd condition of HepG2 cells, with a 4.81-fold increase compared to the NG-UN controls (p < 0.0001; see Fig 6A). In the HUH7 CLEC cell model, superoxide radical production is increased by 3.42-fold in HG-UN, 2.71-fold in HG-1 µM Cd, and 2.84-fold in NG-1 µM, relative to control NG-UN cells. The expected linear increase in superoxide production is seen in HepG2 CLEC cells due to Cd and/or glucose exposures (Fig 6A). However, we note that the HG-1 µM Cd condition of the HUH7 CLEC model showed a decrease in superoxide level (2.71-fold vs 3.42-fold) compared to the hyperglycemia, no Cd exposure (HG-UN; Fig 6B). This non-intuitive reduction in baseline superoxide levels in the HUH7 HG-1 µM Cd cell line may be due to either a. experimental variability or b. due to increased, uncompensated mitochondrial damage in the HUH7 HG-1 µM Cd condition specifically, an issue we explore in later experiments. Similar to the mitochondrial mass, we observe cell-line specific differences in superoxide radical production. Two-way ANOVA analysis detect statistically significant interactions (p < 0.0001) between Cd and glucose effects on superoxide production (see supplementary table 2). Overall, these results establish increased baseline superoxide radical production due to chronic Cd exposures, further amplified by concurrent hyperglycemia. CLEC exposures, in conjunction with hyperglycemia, are thus likely disrupt normal cellular redox homeostasis by driving sustained mitochondrial dysfunction and ROS production. Overall, these results underscore the additive nature of CLEC induced oxidative stress in conjunction with common dysmetabolic states (e.g., type II diabetes) and thus, may play an important role in chronic liver diseases and mitochondrially driven hepatotoxic outcomes.

**Fig. 6.**
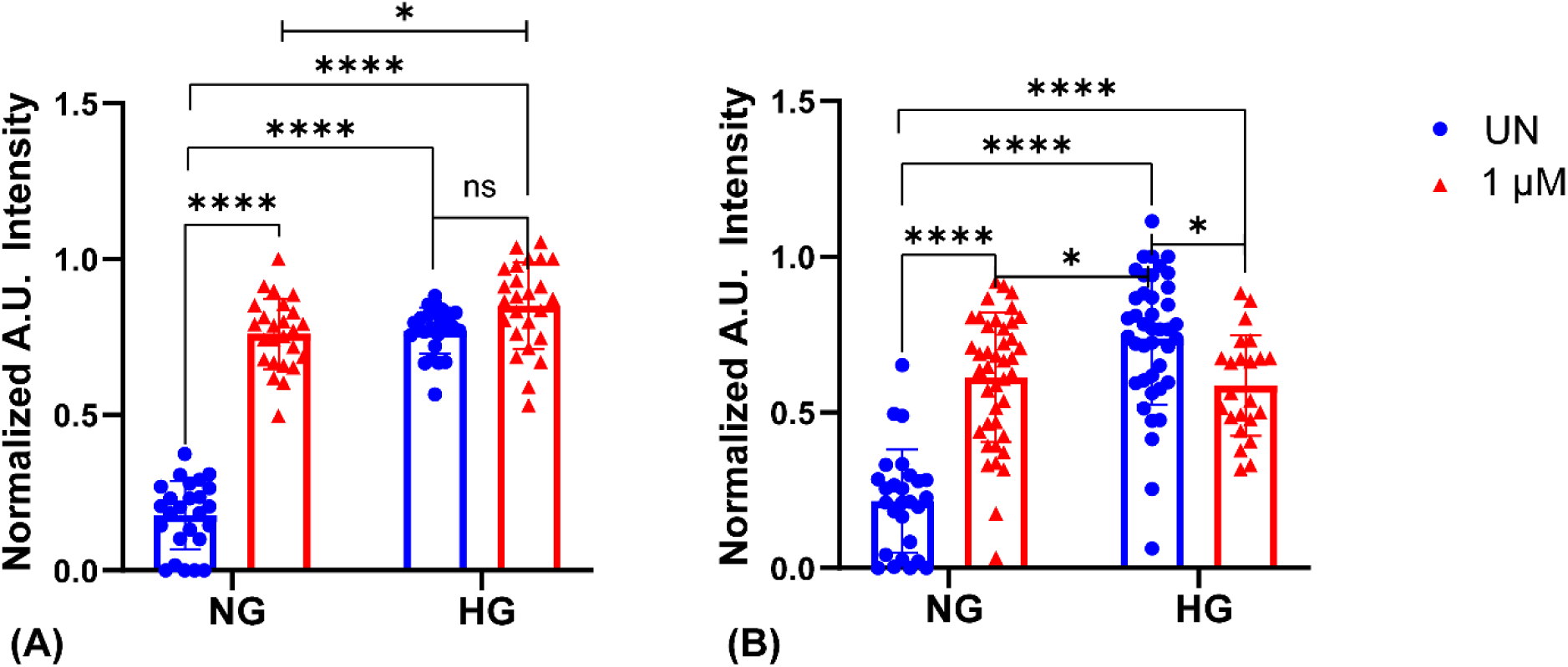
Quantitative measurement of baseline superoxide radical production in CLEC models - The superoxide radical production was measured at baseline in the various CLEC exposure conditions (A) HepG2 and (B) HUH7. Superoxide levels were measured by incubating with 200 nM MitoSox Red dye for a duration of 30 mins. Both CLEC models show elevated levels of superoxide radicals, exacerbated by the hyperglycemia, indicating an additive effect of Cd and hyperglycemia exposures. All the values were normalized using a max - min normalization approach on a 0-1 scale. Data represents the mean of three independent experiments with five well replicates in each plate. Statistical levels of significance are represented as follows - *p<0.05, **p<0.01, ***p<0.001, ****p<0.0001.

### e. Mitochondrial Membrane Potential (MMP) is decreased in CLEC and hyperglycemic cells

To understand the origins of the increased superoxide radical production, we measured the mitochondrial membrane potential (MMP; ΔΨm) of CLEC cells. MMP is the potential gradient across the mitochondrial membrane that is responsible for normal mitochondrial function. Maintaining the MMP gradient is a key barometer of the overall mitochondrial health in all cells including liver cells. Loss of the MMP gradient results in defective ATP production (Kwong & Molkentin, 2015) as well as increased ROS production (Zorov et al., 2006). To understand CLEC and glucose impacts on the MMP, we measured the uptake of an MMP sensitive dye, tetramethyl rhodamine ester (TMRE), whose uptake into the mitochondria is linearly proportional to the MMP gradient (see methods). We observe a monotonic decrease in the TMRE dye fluorescence (NG-UN > NG-1 µM > HG-UN > HG-1 µM) in both HepG2 and HUH7 CLEC cell lines (Fig 7) indicating significant effects of CLEC and glucose exposures on the MMP gradient and an overall decrease in the mitochondrial health. We measured a 2.56-fold decrease in TMRE intensity in the HepG2 CLEC model and a 7.29-fold decrease in HUH7 CLEC between the baseline NG-UN (control) and HG-1 µM (Cd and glucose effects) respectively. Two-way ANOVA analysis confirms statistically significant additive effects (p < 0.0001; see supplementary table 2) of CLEC and glucose exposures on the observed TMRE fluorescence. We also note preferential accumulation of TMRE dye in the perinuclear mitochondria (Fig 7) of Cd exposed cells supporting our prior observations using Mitotracker Green and TOMM20 staining. In summary, sustained CLEC and glucose exposures decrease the MMP gradient driving significant mitochondrial dysfunction which explains the increased superoxide radical production. These findings are of importance in understanding the long-term consequences of Cd exposures and diabetes as drivers of pathological MASLD.

**Fig. 7.**
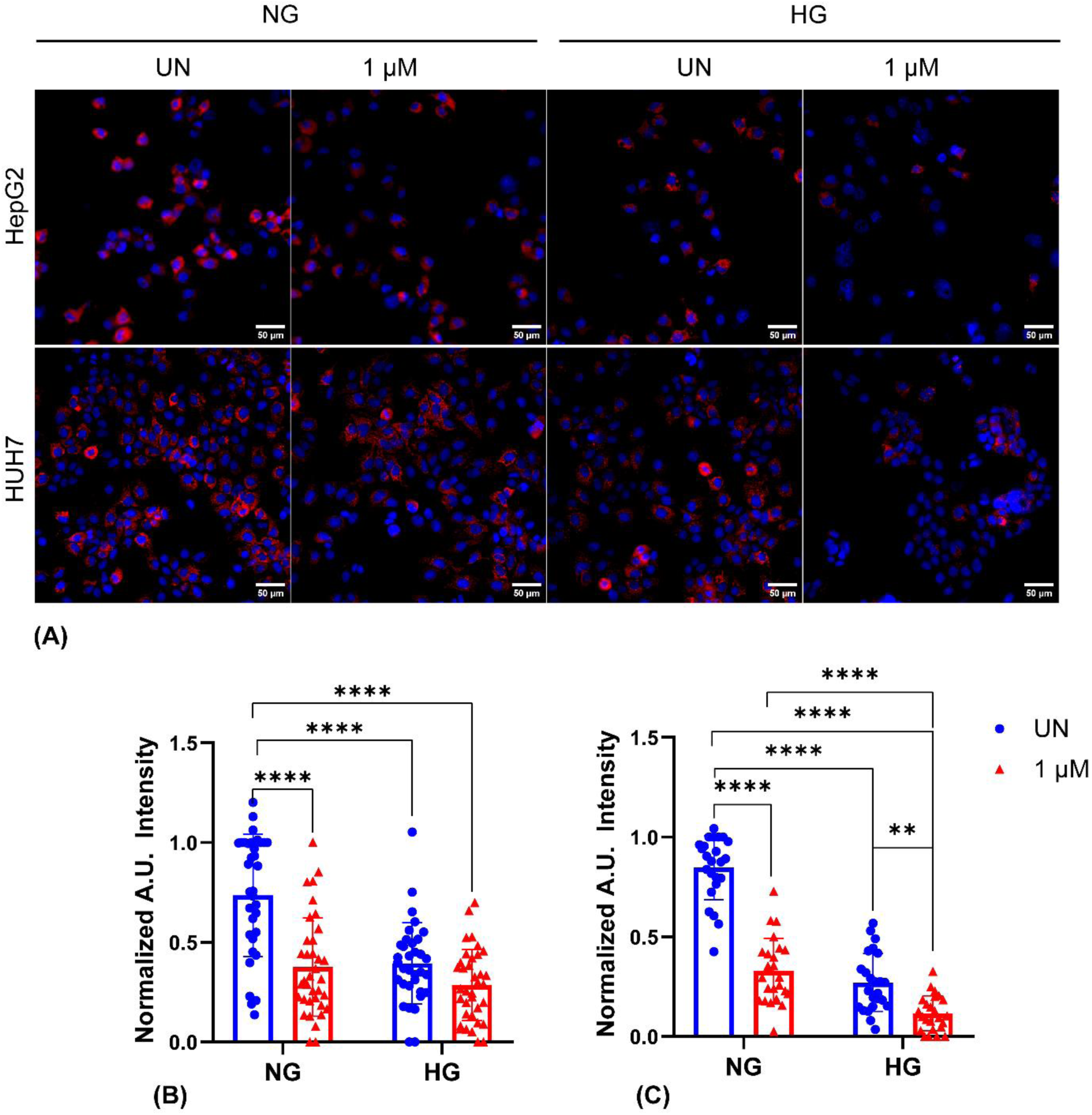
Measurement of mitochondrial membrane potential (MMP; ΔΨm) using TMRE dye. (A) Baseline mitochondrial membrane potential (MMP) was measured in CLEC cells by incubating with 100 nM tetra-methyl rhodamine ester (TMRE) dye for a duration of 30 min. CLEC exposures lead to significant reduction in ΔΨm values across Cd-treated groups in both the cell lines (B) HepG2 and (C) HUH7, which are further exacerbated by superimposed hyperglycemia. All the values were normalized using the max - min normalization method on a 0-1 scale. The data represents a mean of three independent experiments with five well replicates in each plate. The level of significance is identified as follows - *p<0.05, **p<0.01, ***p<0.001, ****p<0.0001.

### **f.** CLEC and hyperglycemia exposures alter baseline oxygen consumption rates (OCRs) while maintaining the spare respiratory capacity

A key unknown is the mitochondrial functionality in cells chronically exposed to heavy metal stress such as Cd with (or without) hyperglycemia. We used the multi-parametric Seahorse assay to measure CLEC, and glucose impacts on mitochondrial functionality. All findings of the Seahorse assay are reported relative to the NG-UN condition which is considered as the baseline to evaluate mitochondrial effects of CLEC and glucose.

Key findings of the assay are summarized from left to right (Fig 8) - i. *Non-mitochondrial respiration* - In both HepG2 and HUH7 CLEC cells, we see significant differences in non-mitochondrial OCRs relative to the NG-UN (Fig 8 a and g), However, a linear trend (↑ or ↓) is not seen with the highest non-mitochondrial OCR values measured in the HG-UN condition. Reduction of non-mitochondrial OCRs in HG-1 µM Cd specifically may be a consequence of direct enzyme inhibition (e.g., NADPH oxidase) due to cytoplasmic free Cd ii. *Basal respiration -* The basal respiration parameter (representing mitochondrial OCR specifically) shows significant differences relative to NG-UN. In the HepG2 CLEC cell line, Cd causes a measurable difference while glucose alone does not. In contrast, in the HUH7 CLEC cell line, a linearly decreasing basal respiratory rate is observed ((NG-UN > NG-1 µM > HG-UN > HG-1 µM; see fig 8 b and h). iii. *ATP-linked respiration* - The ATP-linked respiration parameter (measured by addition of oligomycin – an ATP synthase inhibitor) reveals measurable differences in both HepG2 and HUH7 CLEC cells. However, combined effects of both Cd and glucose on ATP-linked respiration are observed again only in the HUH7 cell line in the form of a linearly decreasing trend (fig 8 c and i). These ATP-specific OCRs findings suggest CLEC and glucose exposures induce differences in this parameter due to either a. ATP utilization changes (e.g., increased/decreased energy demands) b. ATP synthesis differences (direct effect of Cd and/or glucose on ATP synthase) or c. Substrate supply and oxidation differences (e.g., altered MMP values). However, this assay cannot detect the precise source of these ATP-linked respiratory changes. iv. *Maximal respiration* - The maximal respiration due to FCCP induced uncoupling was assessed. While qualitative differences are noted visually (fig d and j), only HG-1µM in HUH7 cells reached statistical significance for measurable differences. This is an unsurprising in light of the need of the cells to maintain a “survivable” reserve of OCRs despite Cd and glucose exposures. Over the 24-week course of the Cd and glucose exposure, the CLEC cell models likely develop compensatory mechanisms (such as increased, if dysfunctional mitochondria) which help the cells cope with an acute maximal respiratory demand due to an inducing insult such as FCCP. v. *Proton leak, and coupling efficiency* - Lastly, we observe that the coupling efficiency (a derived parameter indicating a proportion – i.e., ATP-linked respiration/basal respiration), and the proton leak (fig e, f, k, and l) measures show measurable impacts of CLEC and glucose on HepG2 and HUH7 cells. These derived parameters show differences in both Cd and glucose exposure conditions relative to the NG-UN cells in a cell line specific manner.

**Fig. 8.**
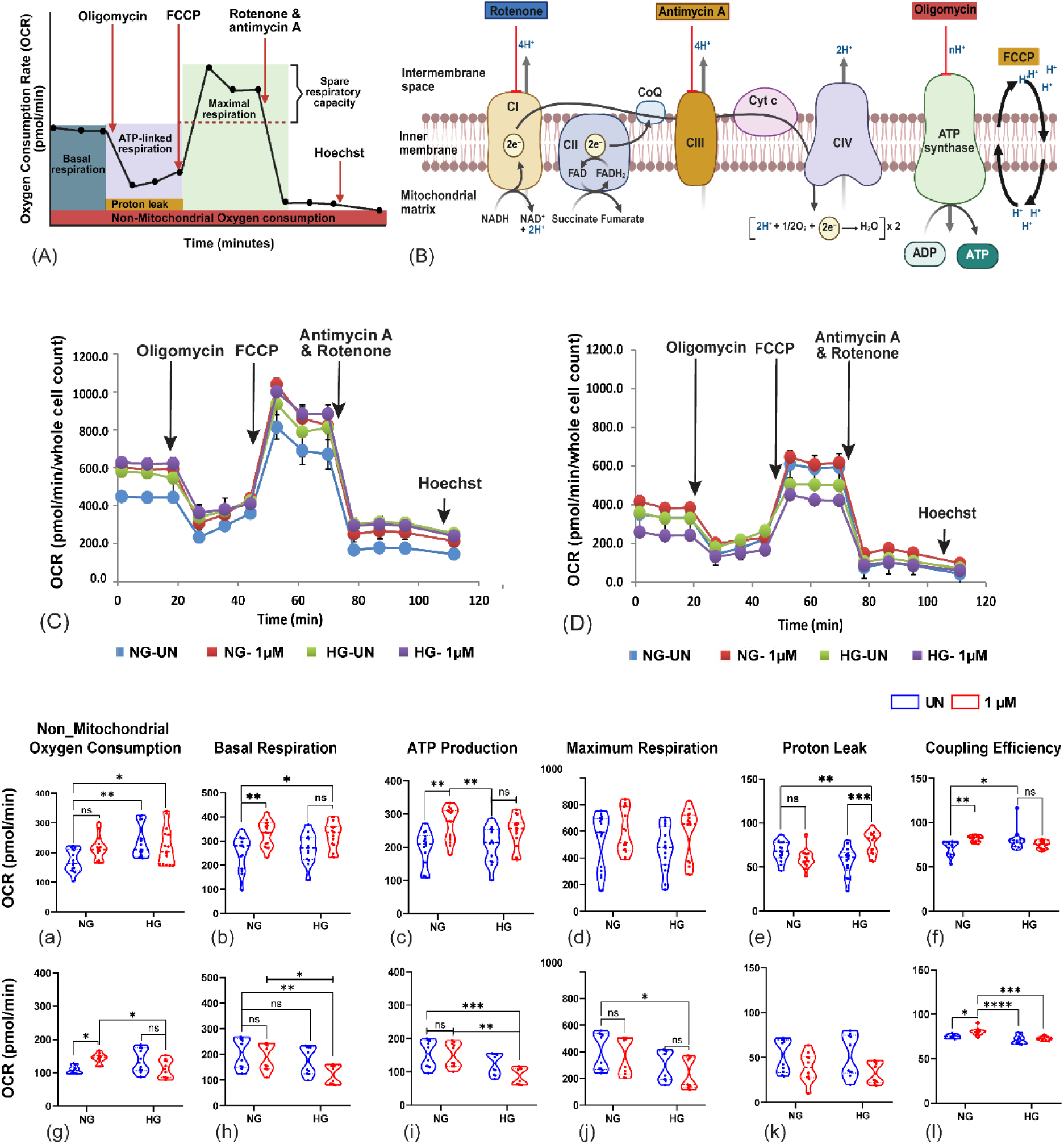
Seahorse MitoStress analysis in the HepG2 and HUH7 CLEC exposure conditions at baseline. (A) Representative data profile of a mitochondrial stress test (OCR) assay (MitoStress Assay). Red arrows denote the time points of the various compound injections in the assay. (B) Schematic indicating the site of action in the mitochondrial electron transport chain (ETC) for the sequential mitochondrial inhibitors injected in each well in the assay: oligomycin, FCCP, rotenone, and antimycin A. (C) The cellular bioenergetic profile (MitoStress) and mitochondrial parameter analysis (a-i) for HepG2 and HUH7 CLEC cell models (C and D respectively). (a – f; top row, g – i; bottom row) Various cellular bioenergetic profile (MitoStress) and mitochondrial parameter outcomes for the HepG2 and HUH7 CLEC models. (top row) HepG2 cells show less alteration of mitochondrial parameters to Cd and hyperglycemia exposures compared to (bottom row) HUH7 CLEC cells which show greater impairment of mitochondrial function. Data values represents the mean of three independent experiments, each with five technical replicates. Statistical significance: The level of significance is represented as *p < 0.05, **p < 0.01, ***p < 0.001, ****p < 0.0001.

In summary, the Seahorse MitoStress assay results establishes CLEC and glucose exposure effects on the baseline OCRs of the mitochondria in both HepG2 and HUH7 cell models. The altered OCRs also likely explain the increased baseline oxidative stress shown previously. Thus, CLEC and glucose exposures likely lead to sustained, low-level damage in the electron transport chains (ETC) of the mitochondria of these cells. However, the mitochondrial damage occurring over a prolonged duration (24-weeks) or dose (1 µM Cd and/or 15 mM glucose) exposures is still not sufficient to cause outright cell death suggesting the presence of progressive compensatory mechanisms of respiration that helps these cells to continue to survive (but not thrive). However, when challenged with an acute mitotoxic insult (e.g., Sorafenib, Oligomycin, FCCP), the CLEC and glucose exposed cells reveal decreased abilities to overcome the acute mitochondrial stress exposure due to the reduced homeostatic mitochondrial respiratory reserves in these cells.

### **g.** CLEC exposure downregulates the antioxidant gene expression

To understand CLEC induced molecular mechanisms of mitotoxicity and explain the differential susceptibility of the cell lines to Cd and glucose (i.e., HUH7 > HepG2), we measured the gene expression of members of key cellular antioxidant pathways. We evaluated the gene expression levels in three main categories: 1. mitochondrial antioxidant genes, 2. cytoplasmic antioxidant genes, 3. genes related to gluconeogenesis pathway (fig 9). The qPCR results show significant downregulation of antioxidant gene expression in HUH7 CLEC cells while HepG2 cells did not show similar changes indicating differential CLEC effects in a cell line specific manner. Significant downregulation of superoxide dismutase 1 (*SOD1*), superoxide dismutase 2 (*SOD2*), catalase (*CAT*), glutathione peroxidase 1 (*GPX1*), glutathione peroxidase 4 (*GPX4*), and NADPH oxidase (*NOX*) is seen in the HUH7 CLEC model cultured under both normoglycemic (5.6 mM) and hyperglycemic (15 mM) CLEC culture conditions. The most prominent downregulation of gene expression was observed in the HG-1 µM Cd condition across multiple genes (1.31 to 3.35-fold range), whereas no such significant changes of gene expression are observed in the HepG2 model (Fig 9).

**Fig. 9.**
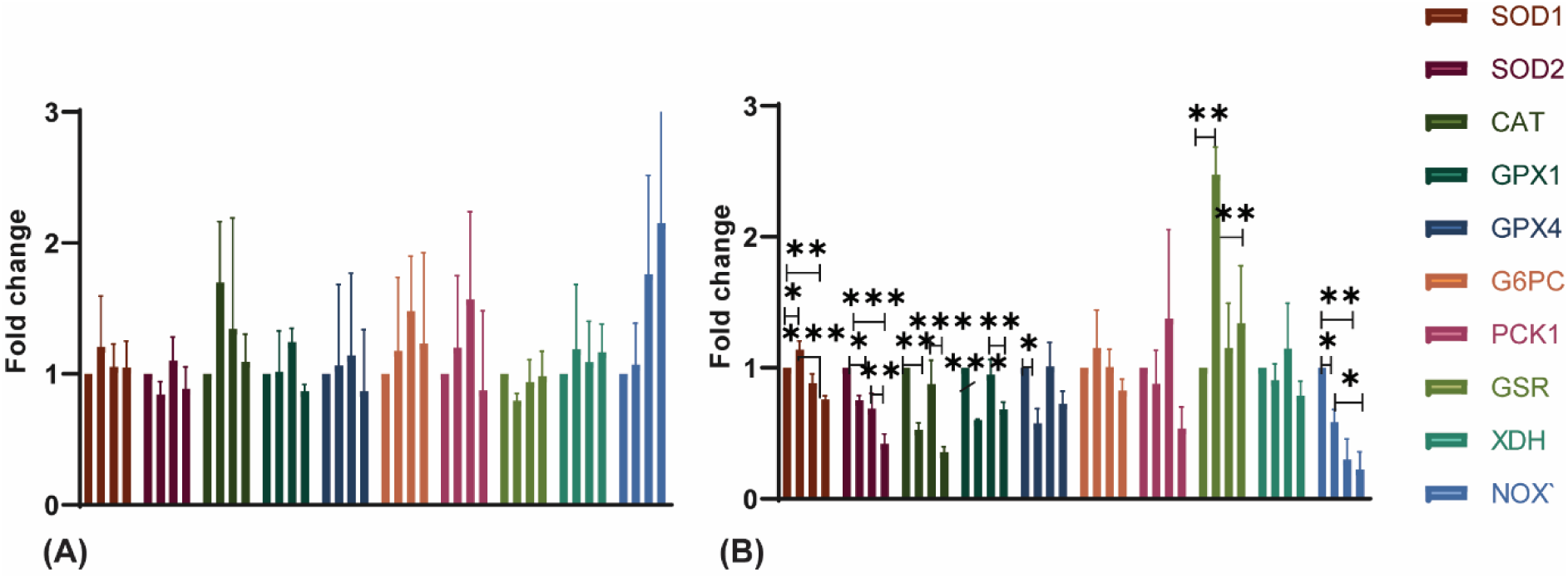
Differential gene expression of anti-oxidant genes in CLEC models – The gene expression of various mitochondrial and cytoplasmic antioxidant genes was measured in (A) HepG2, (B) HUH7 CLEC models. HepG2 cells show a minimal or no significant differences, while HUH-7 cells show statistically significant trends of decreased antioxidant gene expression under CLEC and hyperglycemic conditions. qPCR data represents a mean of three independent biological replicates. Statistical level of significance is represented as follows - *p<0.05, **p<0.01, ***p<0.001, ****p<0.0001.

Three genes - SOD2 (predominantly mitochondrial in expression), CAT (cytoplasmic), and NOX (cytoplasmic) showed the highest downregulation with fold changes of 2.38, 2.82, and 3.35 respectively, in the HG-1 µM Cd condition of the HUH7 CLEC model. The differential cellular gene expression responses seen due to Cd and glucose chronic stress indicate the existence of specific regulatory differences in each cell model. The key finding in this assay establishes CLEC and hyperglycemia exposures as drivers of impaired antioxidant defences, disruptors of cellular redox balance, and promotors of oxidative stress in a context dependent manner. The reduced ability of HUH7 CLEC model (relative to the HepG2) to maintain antioxidant gene expression appears to be directly linked to the difference in the induction of protective metallothionein, an issue explored in the next section. Finally, results from the qPCR assay also show that the defective antioxidant responses associated with mitochondrial dysfunction may occur with hyperglycemic exposures alone (without added Cd), an outcome that is of importance in individuals with metabolic conditions such as type II diabetes.

### **h.** CLEC cells exhibit cell-line specific differences of cellular oxidative stress and gluconeogenesis markers

A key observation thus far, across multiple assays, is the cell line specific impacts of CLEC and glucose exposures. To understand these differences, we measured protein expression of metallothionein (MT-2A) in both cell lines. Metallothionein is a thiol-rich, evolutionarily conserved small protein which plays an important role in sequestering Cd^+2^ ions. We postulated the assay specific differences (HUH7 > HepG2) observed in this study may be due to differences in the induction of the protective metallothionein (MT) gene expression between these cell lines. To establish this, we performed a MT-2A Western blot analysis to support this hypothesis.

We observe a marked induction of MT2A protein expression in CLEC cells, with a fold increase of up to 20.17 seen in HepG2 and 10.48-fold induction seen in HUH7 cells in the HG-1 µM Cd condition relative to NG-UN (see Fig 10 E and F). This is a novel finding in the context of the CLEC hepatocellular model which has not been reported before to our knowledge. We postulate that the relatively high induction of MT-2A seen in the HepG2 CLEC model as compared to HUH7 (20.17-fold vs 10.48-fold) confers a major protective survival advantage by mitigating the free Cd-induced oxidative damage in the HepG2 cell line. In contrast, lowered MT-2A expression as seen in the HUH7 CLEC model may result in increased bioavailability of free Cd^+2^ ions, causing added mitochondrial damage in the HUH7 cell line (explaining the differential assay outcomes observed in this study).

**Fig 10:**
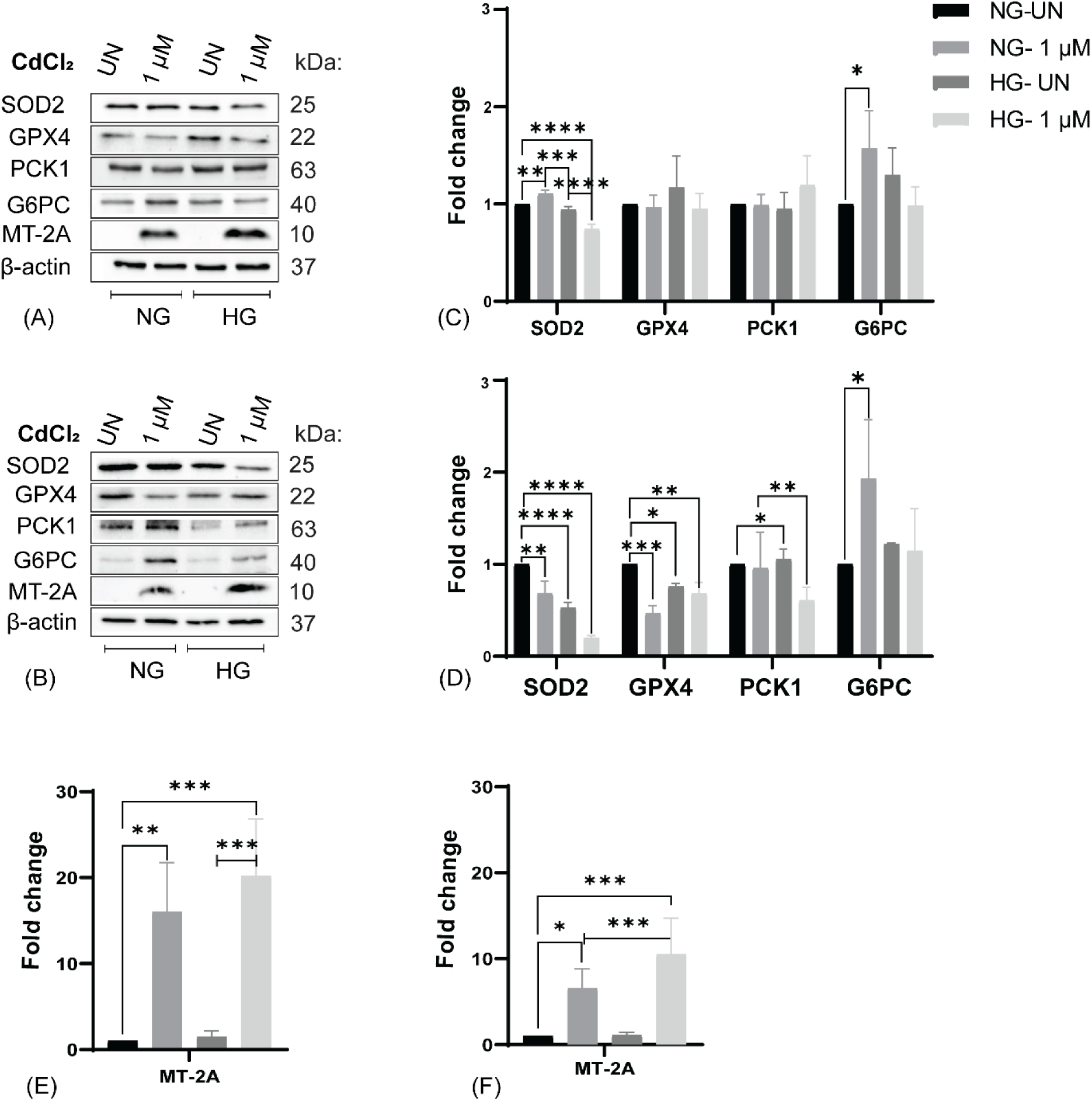
Effects of CLEC and hyperglycemic exposures on antioxidant status and gluconeogenesis – Representative Western blot data - Top (A and C) HepG2 CLEC data. Bottom (B and D) HUH7 CLEC data. Densitometric analysis shows CLEC exposures significantly decreases the SOD2 expression in both CLEC models (C and D) modulated by hyperglycemic (15 mM) exposures. In contrast, GPX4 expression is changed only in the HUH7 CLEC model (D). No effects of CLEC and glucose are noted in the rate-limiting PCK1 and G6PC proteins of the gluconeogenesis pathway (C and D). Significant induction of the metallothionein gene (MT-2A) is observed due to CLEC exposures in both cell lines (Fig 8 E and F) with greater induction in HepG2 compared to HUH7. Also, hyperglycemia induces additional MT-2A expression in both cell lines. All of the densitometric results are reported as the mean ± SEM (n =3). Statistical analyses are performed using two-way ANOVA with Tukey multiple comparison test. The levels of statistical significance are *p*<*0.05, **p*<*0.01, ***p*<*0.001, ****p*<*0.0001.

Increased exposure to free Cd^+2^ ions in the HUH7 CLEC model may also likely result in greater damage to additional proteins seen in mitochondria. To confirm this, we measured protein expression of key proteins of the antioxidant pathway such as superoxide dismutase (SOD2), a manganese dependent, mitochondrial enzyme needed for superoxide radical removal. We measured significant SOD2 expression changes in both the HepG2 and HUH7 CLEC models with the least expression seen in the HG-1 µM Cd model in both cells (Fig 10 A - D) as expected. This is in contrast to expression of glutathione peroxidase 4 (GPX4), a redox scavenging enzyme seen in the cytoplasm, nucleus and mitochondria of cells. Altered GPX4 protein expression reached statistical significance in the HUH7 CLEC model and not the HepG2 CLEC model (fig 10 A - D). Overall, these results indicate increased bioavailability of free Cd^+2^ ions have the potential to affect key antioxidant proteins in the cytoplasm and mitochondria.

Lastly, to understand the hyperglycemia specific effects, we measured the protein expression of glucose-6-phosphatase (G6PC) and phosphoenolpyruvate carboxykinase (PCK) proteins. These are two key metabolic enzymes and the rate-limiting steps of the gluconeogenesis pathway specifically. G6PC and PCK are commonly dysregulated in diabetes (Valera et al., 1994), obesity and cancer (Tan et al., 2025; Yu et al., 2021). However, no significant dysregulation of G6PC was noted due to Cd and glucose exposures.

### **i.** CLEC and hyperglycemia exposures induce increased mitochondrial turnover dynamics in cells

Our initial experiments measuring mitochondrial mass established a decrease due to CLEC and glucose exposures (see results from section c). We confirmed the imaging findings of mitochondrial mass reduction by measuring the TOMM20 protein expression using Western blots. TOMM20 is an integral part of the TOM complex, a key functional unit responsible for the import of proteins into the mitochondria. We find CLEC and glucose dependent reduction of the TOMM20 protein expression in both HepG2 and HUH7 cell lines confirming the initial imaging findings of Mitotracker Green and confocal TOMM20 staining. A 1.8-fold and 2.94-fold reduction of TOMM20 expression is seen in the HG-1 µM Cd condition relative to the NG-UN condition in HepG2 and HUH7 respectively (see fig 11 A – D).

**Fig 11:**
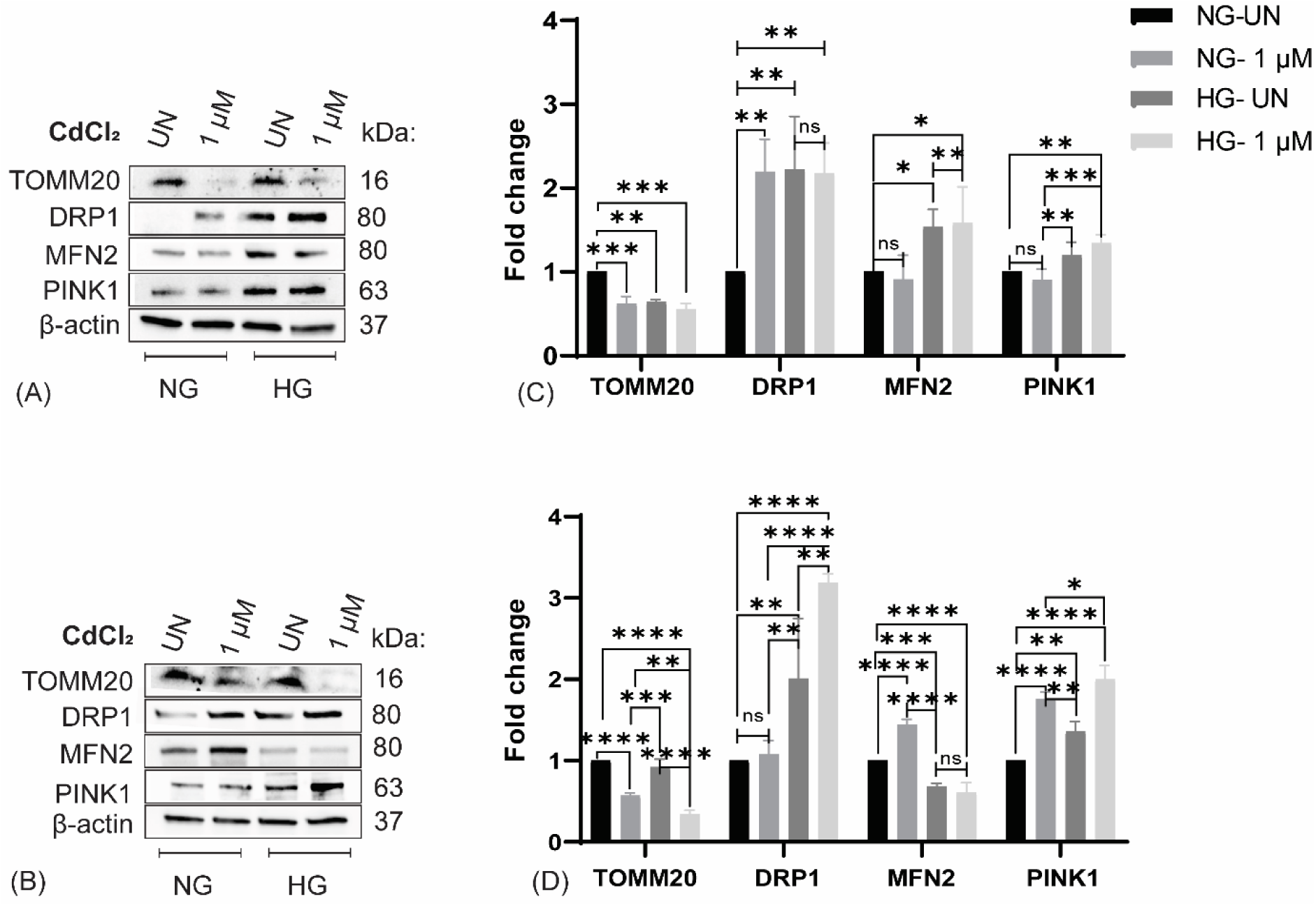
Effects of CLEC and hyperglycemia on mitochondrial morphology and dynamics – Representative Western blot data - Top (A) HepG2 CLEC data. Bottom (B) HUH7 CLEC data. Densitometric analysis shows CLEC exposures significantly decrease the TOMM20 expression in CLEC models (C and D) which is further modulated by concurrent hyperglycemic (15 mM) exposures. CLEC models (HepG2 and HUH7) show increased DRP1 gene expression (Fig 11 C and D) driving increased mitochondrial fission in CLEC cells. Mitofusin (MFN2) protein shows opposite trends with increase in HepG2 (Fig 11 C) while decrease in HUH7 (Fig 11 D). Increase PINK1 expression is seen in both cell lines indicating increased mitochondrial turnover due to Cd and hyperglycemia. All densitometric results are reported as the mean ± SEM (n =3). Statistical analyses are performed using two-way ANOVA with Tukey multiple comparison test. The levels of statistical significance are *p*<*0.05, **p*<*0.01, ***p*<*0.001, ****p*<*0.0001.

To understand the specific impacts of CLEC and hyperglycemia on mitochondrial biogenesis, we measured dynamin-related protein 1 (DRP1), a key protein involved in the regulation of fission of cellular mitochondria. Prior studies showed acute high-dose Cd exposures induce DRP1 expression (Xu et al., 2013). Similarly, we observe an increased expression of DRP1 protein due to CLEC and hyperglycemia exposures in this study (see fig 11 A-D). This is particularly striking in the HUH7 CLEC model which shows a 3.18-fold increase of DRP1 expression (HG-1µM Cd vs NG-UN baseline). In the HepG2 CLEC model, a 2.2-fold induction of DRP1 expression is observed (HG-1µM Cd vs NG-UN baseline; Fig B and D). These findings confirm altered mitochondrial biogenesis/dynamic effects in response to the long-term, CLEC and glucose exposures as well (as opposed to acute Cd exposures alone).

The protein expression of mitofusion 2 (MFN2), a key member of the MFN1/2 complex of proteins responsible for mitochondrial fusion (instead of fission) was measured. While the HUH7 CLEC model showed an anticipated decreasing trend of MFN2 expression with HG-UN and HG-1 µM Cd, the HepG2 CLEC model showed an opposite trend (↑ MFN2 expression in HG-UN and HG-1uM Cd; Fig A -D) which may be due to cell line variability. Lastly, to assess the impacts of CLEC and glucose on mitochondrial quality control in response to environmental stressors (Cd and glucose), we measured the expression of PTEN-induced kinase 1 (PINK1), a central regulator of mitophagic processes. We measure increased PINK1 expression in both HepG2 and HUH7 CLEC cells, with the highest induction of PINK1 in the HG-1µM Cd condition (1.5-fold and 2-fold respectively).

Elevated PINK1 expression indicates activation of the mitophagic pathway, likely in an attempt to actively remove the damaged mitochondria due to CLEC and hyperglycemic stress and ensure mitochondrial quality control. The differences in the DRP1, MFN2 and PINK1 induction seen in our model suggests complex (and dynamic) regulatory effects in response to environmental exposures such as Cd and glucose and requires further study.

## IV. Discussion

Chronic, heavy metal-induced cellular stress is increasingly recognized as a key, but underappreciated, factor driving metabolic disease etiopathology (e.g., type II diabetes; T2DM) and hepatocellular dysfunction (Haidar et al., 2023). Although Cd-induced nephrotoxicity has been extensively studied (Johri et al., 2010; Ma et al., 2022), the long-term consequences of Cd exposure on hepatic mitochondrial health remains poorly understood (Cannino et al., 2009). Multiple epidemiological and experimental studies (both acute and chronic) establish an association of Cd exposures with the onset (and progression) of metabolic syndrome, T2DM, and metabolic (dysfunction) associated fatty liver disease (MASLD) (Buha et al., 2020; Li et al., 2023; Wang et al., 2018). Despite these observations, relatively few toxicological studies exist reviewing the role of *chronic* heavy metal exposures in the context of pre-existing metabolic dysfunction, such as type II diabetes (Buha et al., 2020; Kuo et al., 2013; Sun et al., 2022). To address this gap, we conducted a novel *in vitro* dual-exposure model that combines physiologically relevant chronic low-dose Cd exposures (CLEC) with a high glucose (15 mM) T2DM mimicking condition to understand their impact on mitochondrial function in this study. Unlike prior studies that examine either Cd toxicity (Korotkov, 2023; Odewumi et al., 2018) or hyperglycemia separately (Wang et al., 2022), the current study implements a new approach method (NAMs) to generate key mechanistic toxicological insights into the combined impacts of Cd and glucose on mitochondrial function and hepatocellular homeostasis.

This study establishes hepatic cells with simultaneous (and chronic) Cd and glucose exposures show significant cytotoxic and mitotoxic outcomes. At baseline, EC_50_ values for Sorafenib exposure are approximately similar in all conditions, suggesting glycolytic compensation over the long duration of the Cd and glucose exposure. This observation aligns with the prior studies which showed that immortalized cell lines with enhanced glycolytic flux can transiently tolerate mitochondrial insults (Oldani et al., 2020; Xu et al., 2005; Yu et al., 2017), because high glucose availability favours glycolysis as the primary ATP source for homeostasis (Kukurugya et al., 2024). However, when switched to galactose-containing media, this enforces selective reliance on mitochondrial oxidative phosphorylation (Skolik et al., 2021). The protective compensatory effect of glycolysis is thus abolished, unmasking the true extent of mitochondrial damage due to long-term CLEC and glucose exposures. From a clinical perspective, these findings raise concern that compared to normoglycemic individuals, hyperglycemic diabetic patients with concomitant environmental Cd exposures may face heightened risks of liver injury (due to lowered homeostatic thresholds) in response to common day-to-day episodic drug or toxin exposures. This concept is consistent with prior epidemiological studies linking Cd exposures to T2DM and liver disease progression (Park et al., 2021; Schwartz et al., 2003). These findings also highlight the need to evaluate drug toxicity outcomes in the context of pre-existing metabolic and environmental stressors rather than assuming baseline patient uniformity for characterizing hepatotoxic outcomes and predictors of liver injury (Le Magueresse-Battistoni et al., 2018; Sun et al., 2022).

An important factor of mitochondrial redox homeostatic balance is maintenance of the mitochondrial membrane potential, a fundamental driver of cellular energy production. Several in-vivo and in-vitro studies show the effect of acute Cd exposures directly compromise cellular mitochondrial function by altered GSH/GSSG redox balance, altered mitochondrial dynamics and inhibition of mitochondrial respiration (Belyaeva et al., 2006; Belyaeva et al., 2004). It has been reported that Cd exposures cause a direct collapse of mitochondrial membrane potential in primary hepatocytes (Liu & Liun, 1990; Martel et al., 1990), and cancer cell lines (Li et al., 2000; Zhang et al., 2008) causing increased oxidative stress. However, most of these studies examined the effects of acute cadmium exposures. In the current chronic heavy metal stress focused study, we observed the least MMP gradient in the HG-1 µM Cd CLEC condition, indicating the cell mitochondria are unable to maintain the MMP gradient due to CLEC and glucose exposures. In addition to the chronic MMP dysfunction (depolarization), changes are noted in the mitochondrial mass, morphology and the increased mitochondrial superoxide production of both the CLEC cell models in this study.

Direct measurement of mitochondrial respiratory dysfunction is a sensitive marker of mitochondrial toxicity (Hynes et al., 2006; Swiss et al., 2013). Cadmium inhibits the activity of mitochondrial respiratory complexes in hepatocytes (Belyaeva et al., 2011; Muller & Ohnesorge, 1984). Such inhibition disrupts mitochondrial function, compromising the cell’s ability to meet increased energy demands and thereby reducing the mitochondrial parameters such as basal respiration, ATP production, and reserve capacity, an indicator of bioenergetic flexibility (Desler et al., 2012). Functional mitochondrial damage is characterized by decreased mitochondrial membrane potential, elevated superoxide production, enhanced lipid peroxidation, and depletion of endogenous antioxidant defenses (Lolescu et al., 2025; Xu et al., 2025). Cd^2+^ has been shown to increase superoxide radical production directly, by targeting proteins in the complexes I, III, and IV of the electron transport chain (Belyaeva et al., 2011), and disrupted calcium homeostasis (Zhou et al., 2015). Cd^2+^ exposures may also affect mitochondrial function indirectly, by depleting antioxidant defences such as glutathione (Nigam et al., 1999; Taysi, 2024). Hyperglycaemia exacerbates these effects by driving mitochondrial substrate excess, leading to increased reduction of the electron carriers and excess electron flux and leaking of electrons from ETC pathway, causing premature oxygen reduction to superoxide radicals (Yan, 2014).

All of these findings are indeed seen within this study, strongly suggesting that CLEC and hyperglycemia additively perturb the electron transport cycle and tip the redox balance of liver cells towards a state of chronic oxidative stress. It is noteworthy that the CLEC cell models in this study do not show elevated levels of apoptosis at the end of the exposure period, indicating the existence of cellular compensatory/protective mechanisms over prolonged durations of Cd and glucose exposures. Thus, sustained low-dose heavy metal stress (such as Cd) may result in functional cell compromise without active cell replacement via cell death. This finding is important to understand the pathophysiology of diabetes-related diseases which show impaired respiratory capacity, altered mitochondrial ultrastructure, and dysregulated expression of respiratory chain complexes of cells (Verma et al., 2017). In the current study, the Seahorse XF analysis similarly confirms impaired mitochondrial respiration and reduced bioenergetic capacities due to CLEC and glucose exposures in hepatic cells which would be of relevance to understand long-term metabolically driven hepatic diseases such as hepatic insulin resistance and MASLD.

Since mitochondrial respiratory capacity is tightly coupled to the cellular redox health of cells, we evaluated gene expression of key antioxidant genes in the redox pathway. In spite of the elevated oxidative stress, we observed decreased expression in multiple genes of the antioxidant pathway which was puzzling. Furthermore, our Western blot assay data confirmed the qPCR gene expression findings via reduced expression of MnSOD (SOD2; a key mitochondrial antioxidant gene) seen in both CLEC cell models. Considering the prolonged duration of exposure implemented in this study (i.e., 24 weeks), the reduced antioxidant gene expression observed likely reflects long-term damage processes due to CLEC and hyperglycemia exposures. Heavy metal exposures are associated with long-term cell damaging effects such as advanced glycation end-product (AGE) accumulation (Huo et al., 2023), and genomic or epigenetic modifications (e.g., repressive histone marks on the *SOD2* promoter) (Liu et al., 2024). Chronic hyperglycemia is well-established to induce non-enzymatic glycation (Khalid et al., 2022) in diabetic individuals, along with the nitration of antioxidant proteins, which decreases their enzymatic efficiency over time (Elshaer et al., 2018; Stavniichuk et al., 2014). Such chronic cell damaging effects represent a key element of human diabetic pathophysiology. Thus, the dysregulation of gene and protein-level expression of the antioxidant pathway genes observed in this study may indicate compromised antioxidant capacity due to the sustained oxidative cell damage. Reduced antioxidant function may drive compromised hepatic liver function over prolonged CLEC and glucose exposures and requires further studies in different models.

A major finding of this study is the impact of CLEC and hyperglycemic exposures have on cellular mitochondrial dynamics. In this study, we observe the transition of mitochondria from an elongated, filamentous mitochondrial normal network appearance to a fragmented, edematous and disrupted mitochondrial network appearance seen in the CLEC exposure models. Additionally, we observe perinuclear accumulation of the mitochondria in the CLEC conditions for the first time. These morphological alterations due to long-term CLEC and glucose exposures in this study are consistent with the mitochondrial fragmentation, impaired total mitochondrial mass (Wen et al., 2022) and compromised organelle function reported in other studies due to environmental exposures (Perdiz et al., 2019). Disruption of mitochondrial morphology and dynamics provide the key mechanistic link between chronic heavy metal exposure stress and the long-term impairment of hepatocellular function due to loss of mitochondrial function such as increased mitochondrial permeability transitions, decreased respiratory chain activity, decreased ATP production, and induction of oxidative stress that have been reported in previous studies of MASLD etiopathogenesis (Lolescu et al., 2025; Zhong et al., 2025).

Acute Cd exposures have been shown to disrupt the homeostatic balance of mitochondria by activating a mitochondrial fission mediator, dynamin-related protein 1 (DRP1), often via Ca²⁺ signaling dysregulation (Xu et al., 2013). Mitochondrial calcium homeostatic dysfunction is well recognized as a key mechanism underpinning Cd toxicity, attributable to Cd’s ability to mimic essential divalent cations such as Ca²⁺ and Zn²⁺, displacing them via stronger binding in proteins and disrupting their essential biochemical role in cells. Similarly, hyperglycemia has been shown to promote excessive fission and altered mitochondrial morphologies (Kumari et al., 2012; Yu et al., 2006). Mitochondria are highly dynamic organelles whose functional integrity depends on the dynamic balance between mitochondrial fusion and fission occurring inside the cells. Disruption of this equilibrium is implicated in the pathogenesis of diverse diseases, including neurodegenerative disorders, cardiovascular pathologies, and diabetes mellitus (Adebayo et al., 2021; Geto et al., 2020).

In this study, we measured a significant increase in dynamin-related protein 1 (DRP1) expression, a master regulator of mitochondrial fission. This molecular signature aligns with earlier studies reporting that CdCl₂ triggers DRP1-dependent mitochondrial fission, while silencing or pharmacological inhibition of DRP1 effectively rescued mitochondrial function, while reducing the Cd-induced cytotoxicity (Xu et al., 2013; Zhang et al., 2019). Manipulation of DRP1 activity in Cd-injured rat livers has been shown to mitigate mitochondrial fragmentation and preserve organelle function. In this study, maximal DRP1 expression (and mitochondrial fission) is seen in the HG + 1µM Cd condition, suggesting DRP1-driven mitochondrial fragmentation is a key driver of the mitochondrial dysfunction in our CLEC model. Fragmented mitochondria not only exhibit reduced ΔΨm, but also reduced overall bioenergetic efficiency while generating increased superoxide radicals, driving the chronic oxidative stress condition of cells. These findings are of key importance in establishing the role of environmental heavy metal exposures as a key driver of mitochondrial dysfunction and MASLD pathogenesis.

In addition to dysfunction of the mitochondrial dynamics, we also observe increased expression of the mitophagy relevant PINK1 protein in both CLEC models. PINK1 accumulation on damaged mitochondria is a key initiating event of mitophagy necessary for cell maintenance and survival (Wang et al., 2023). Elevated PINK1 levels seen in the CLEC models indicate active mitophagic pathways, removing the damaged mitochondria due to CLEC and glucose stress. However, while such mitophagic clearance may transiently preserve cellular viability, over the long term, persistent mitophagic activation can cause significant mitochondrial depletion (Wang et al., 2023) and impaired respiratory reserves (Aravamudan et al., 2017; Prakash et al., 2017) reinforcing the hepatic bioenergetic dysfunction loop that is commonly seen in MASLD. Our findings therefore strengthen the emerging consensus that excessive mitochondrial fragmentation not only precedes but is also a necessary step for Cd-induced mitochondrial dysfunction in cells.

Over the course of this study, we identified elevated impacts of CLEC and glucose exposures in multiple assays (cell viability, superoxide production, MMP, Seahorse experiments, qPCR, and Western blots) in the HUH7 CLEC model (compared to the HepG2) which was puzzling. We speculated these minor, but definite, differences may be due to cell-line specific effects of Cd and glucose. To validate this hypothesis, we measured metallothionein, an evolutionarily conserved Cd-binding protein that mitigates heavy metal redox stress. We measured significant differences in the induction of the specific MT2A protein in HepG2 (+++) compared to HUH7 (++) which supported our hypothesis. Indeed, literature exists which establishes associations of metallothionein protein polymorphisms to variable disease and cancer susceptibilities among different individuals. Metallothionein (MT) gene polymorphisms (e.g., frequently studied rs28366003 and rs10636 polymorphisms) in the MT2A gene are linked to increased risk of breast (Liu et al., 2017), prostate (Forma et al., 2012), and laryngeal cancers (Starska et al., 2014). MT gene polymorphisms can affect detoxification, antioxidant defence, and cellular processes like proliferation and differentiation. The specific role of MT polymorphisms in the context of chronic liver diseases such as MASLD needs further study and validation. Thus, our current CLEC model underscores the importance of understanding the molecular and cellular context while measuring mitotoxicity and/or therapeutic responses. More broadly, these findings highlight the broader issue of the role of human genetic variation in our ability to sequester heavy metals effectively. Humans likely respond in a variable manner to environmental pollutants which may be of importance in the study of a complex disease such as MASLD.

Overall, this study illustrates key important themes of MASLD pathogenesis – a. the important role played by chronic heavy metal stress (CLEC; an external factor) and glucose levels (an internal factor) as drivers of metabolic (and mitochondrial) dysfunction seen in liver cells, b. the non-trivial impacts of pollutants on mitochondrial and cellular homeostasis, even at “low-doses” when they occur over extended periods of time and c. hyperglycemia as a significant source of mitochondrial dysfunction and increased oxidative stress in liver cells. Data from this study supports the emerging paradigm that chronic mitochondrial dysfunction (regardless of proximal cause) can drive hepatic metabolic disease and environmental toxicant-induced hepatocellular injury. Additionally, preserving (and recovering) mitochondrial function in response to environmental toxic injury may be an important therapeutic MASLD strategy, while focusing on preventing environmentally induced hepatic mitotoxicity. In summary, the current study leverages a novel invitro hepatocellular model to establish the role of chronic, low-dose Cd (CLEC) exposures and hyperglycemia as a driver of chronic mitochondrial function.

## V. Conclusions

The vast majority of invitro studies in toxicology focus on acute, high-dose exposures representing non-physiological exposure models of toxic heavy metals pollutants. Yet, to understand the true pathophysiological outcomes of environmental heavy metal pollution in the real-world context, novel chronic low-dose models such as the current study are needed. In this study, we examined the impacts of chronic, low-dose exposures of Cd along with normo- and hyperglycemia conditions, thus modelling realistic outcomes. We established the critical role of CLEC and glucose exposures as drivers of hepatic mitochondrial dysfunction. This study also establishes the significant additive effects of cadmium and glucose exposures as drivers of long-term hepatic cellular and mitochondrial dysfunction. Outcomes measured in this study help us to gain a focused perspective on the key mitochondrial dysfunction that may be driving the observed global increases of metabolic diseases such as type II diabetes, obesity, and MASLD.

### CRediT authorship contribution statement

Rahul Kumar: Writing – original draft, data collection, data analysis, data visualization, and editing. Ashwin Chinala: Writing – draft revision and editing. Li Chen: Writing – review and feedback. Sharina Desai – review and feedback. Marcus Garcia – review and feedback. Sarah N. Blossom – review and feedback. Matthew Campen – review and feedback. Rama R. Gullapalli: Writing – original draft, editing, visualization, supervision, funding acquisition, and conceptualization.

## Funding Acknowledgements

Authors were supported by a grant from the UNM Center for Metals in Biology and Medicine (CMBM) through NIH NIGMS grant P20 GM130422. RRG was supported by a pilot award from the NM-INSPIRES P30 grant 1P30ES032755. This work was supported by AIM centre core funded by NIH grant P20GM121176

---

**Supplementary table 1.**
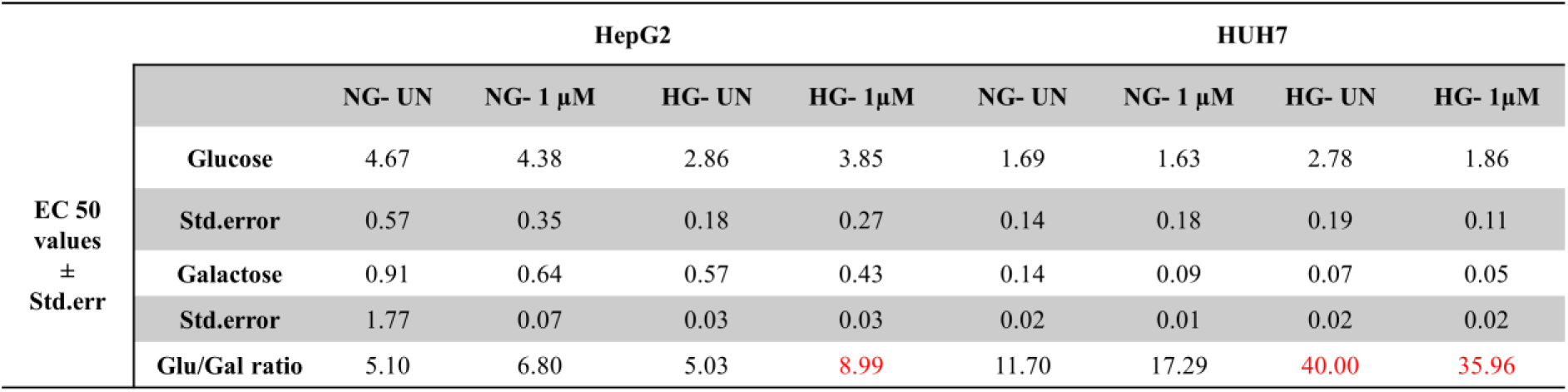
EC50 values and measured Glu/Gal ratios.

**Supplementary table 2.** Two-way ANOVA analysis table.

A. Mitochondrial Mass Data
| Cell line | Glucose concentration<br>(5.6mM, 15 mM) | Cadmium concentration<br>(0 nM, 200 nM, 1 µM) | Interaction<br>(Glucose * cadmium) |
| --- | --- | --- | --- |
| HepG2 | 0.0226 | <0.0001 | 0.0003 |
| HUH7 | <0.0001 | <0.0001 | 0.0811 |

B. MitoSox (Superoxide) Radical
| Cell line | Glucose concentration<br>(5.6mM, 15 mM) | Cadmium concentration<br>(0 nM, 200 nM, 1 µM) | Interaction<br>(Glucose * cadmium) |
| --- | --- | --- | --- |
| HepG2 | <0.0001 | <0.0001 | <0.0001 |
| HUH7 | 0.0020 | <0.0001 | <0.0001 |

C. Mitochondrial Membrane Potential (MMP)
| Cell line | Glucose concentration<br>(5.6mM, 15 mM) | Cadmium concentration<br>(0 nM, 200 nM, 1 µM) | Interaction<br>(Glucose * cadmium) |
| --- | --- | --- | --- |
| HepG2 | <0.0001 | <0.0001 | 0.0023 |
| HUH7 | <0.0001 | <0.0001 | <0.0001 |

D. Seahorse data i. Non-Mitochondrial Basal Respiration
| Cell line | Glucose concentration<br>(5.6mM, 15 mM) | Cadmium concentration (0<br>nM, 200 nM, 1 µM) | Interaction<br>(Glucose * cadmium) |
| --- | --- | --- | --- |
| HepG2 | 0.0084 | 0.30115 | 0.0278 |
| HUH7 | 0.8375 | 0.5439 | 0.0009 |

ii. Basal Respiration
| Cell line | Glucose concentration<br>(5.6mM, 15 mM) | Cadmium concentration<br>(0 nM, 200 nM, 1 µM) | Interaction<br>(Glucose * cadmium) |
| --- | --- | --- | --- |
| HepG2 | 0.7389 | 0.0002 | 0.5330 |
| HUH7 | 0.0045 | 0.0386 | 0.2287 |

iii. Maximal Respiration
| Cell line | Glucose concentration<br>(5.6mM, 15 mM) | Cadmium concentration (0<br>nM, 200 nM, 1 µM) | Interaction<br>(Glucose * cadmium) |
| --- | --- | --- | --- |
| HepG2 | 0.4582 | 0.0839 | 0.6064 |
| HUH7 | 0.0087 | 0.3350 | 0.9118 |

iv. Proton Leak
| Cell line | Glucose concentration<br>(5.6mM, 15 mM) | Cadmium concentration<br>(0 nM, 200 nM, 1 µM) | Interaction<br>(Glucose * cadmium) |
| --- | --- | --- | --- |
| HepG2 | 0.2805 | 0.0877 | <0.0001 |
| HUH7 | 0.6294 | 0.0081 | 0.6033 |

v. ATP Production
| Cell line | Glucose concentration<br>(5.6mM, 15 mM) | Cadmium concentration (0<br>nM, 200 nM, 1 µM) | Interaction<br>(Glucose * cadmium) |
| --- | --- | --- | --- |
| HepG2 | 0.6314 | 0.0001 | 0.1844 |
| HUH7 | 0.0001 | 0.0886 | 0.1258 |

vi. Coupling Efficiency
| Cell line | Glucose concentration<br>(5.6mM, 15 mM) | Cadmium concentration<br>(0 nM, 200 nM, 1 µM) | Interaction<br>(Glucose * cadmium) |
| --- | --- | --- | --- |
| HepG2 | 0.6423 | 0.1338 | 0.0003 |
| HUH7 | <0.0001 | 0.0080 | 0.1035 |

E. Western Blot Densitometric Analysis i. SOD2
| Cell line | Glucose concentration<br>(5.6mM, 15 mM) | Cadmium concentration (0<br>nM, 200 nM, 1 µM) | Interaction<br>(Glucose * cadmium) |
| --- | --- | --- | --- |
| HepG2 | <0.0001 | 0.0140 | <0.0001 |
| HUH7 | 0.0002 | <0.0001 | 0.8140 |

ii. GPX4
| Cell line | Glucose concentration<br>(5.6mM, 15 mM) | Cadmium concentration<br>(0 nM, 200 nM, 1 µM) | Interaction<br>(Glucose * cadmium) |
| --- | --- | --- | --- |
| HepG2 | 0.4094 | 0.1976 | 0.3384 |
| HUH7 | 0.7614 | 0.0001 | 0.0007 |

iii. MT-2A
| Cell line | Glucose concentration<br>(5.6mM, 15 mM) | Cadmium concentration (0<br>nM, 200 nM, 1 µM) | Interaction<br>(Glucose * cadmium) |
| --- | --- | --- | --- |
| HepG2 | 0.3125 | <0.0001 | 0.4189 |
| HUH7 | 0.1141 | <0.0001 | 0.1321 |

iv. G6PC
| Cell line | Glucose concentration<br>(5.6mM, 15 mM) | Cadmium concentration<br>(0 nM, 200 nM, 1 µM) | Interaction<br>(Glucose * cadmium) |
| --- | --- | --- | --- |
| HepG2 | 0.3088 | 0.3636 | 0.0079 |
| HUH7 | 0.1926 | 0.0591 | 0.0312 |

v. PCK1
| Cell line | Glucose concentration<br>(5.6mM, 15 mM) | Cadmium concentration (0<br>nM, 200 nM, 1 µM) | Interaction<br>(Glucose * cadmium) |
| --- | --- | --- | --- |
| HepG2 | 0.3829 | 0.2155 | 0.1815 |
| HUH7 | 0.0282 | 0.0952 | 0.0282 |

vi. TOMM20
| Cell line | Glucose concentration<br>(5.6mM, 15 mM) | Cadmium concentration (0<br>nM, 200 nM, 1 µM) | Interaction<br>(Glucose * cadmium) |
| --- | --- | --- | --- |
| HepG2 | 0.0007 | 0.0004 | 0.0054 |
| HUH7 | 0.0019 | <0.0001 | 0.0659 |

vii. DRP1
| Cell line | Glucose concentration<br>(5.6mM, 15 mM) | Cadmium concentration<br>(0 nM, 200 nM, 1 µM) | Interaction<br>(Glucose * cadmium) |
| --- | --- | --- | --- |
| HepG2 | 0.0129 | 0.0165 | 0.0113 |
| HUH7 | <0.0001 | 0.0204 | 0.0401 |

viii. MFN2
| Cell line | Glucose concentration<br>(5.6mM, 15 mM) | Cadmium concentration (0<br>nM, 200 nM, 1 µM) | Interaction<br>(Glucose * cadmium) |
| --- | --- | --- | --- |
| HepG2 | 0.8509 | 0.0002 | 0.5911 |
| HUH7 | <0.0001 | 0.0003 | <0.0001 |

ix. PINK1
| Cell line | Glucose concentration<br>(5.6mM, 15 mM) | Cadmium concentration<br>(0 nM, 200 nM, 1 µM) | Interaction<br>(Glucose * cadmium) |
| --- | --- | --- | --- |
| HepG2 | <0.0001 | 0.9760 | 0.1148 |
| HUH7 | 0.0015 | 0.0595 | 0.5697 |

